# A Comprehensive Roadmap of Human Placental Development *in vitro*

**DOI:** 10.1101/2022.04.07.487558

**Authors:** Jaroslav Slamecka, Carlos A. Tristan, Seungmi Ryu, Pei-Hsuan Chu, Claire Weber, Tao Deng, Yeliz Gedik, Pinar Ormanoglu, Sam Michael, Ty C. Voss, Anton Simeonov, Ilyas Singeç

**Author notes:** Correspondence: Dr. Ilyas Singeç, NIH National Center for Advancing Translational Sciences (NCATS) Stem Cell Translation Laboratory (SCTL), NIH Regenerative Medicine Program 9800 Medical Center Drive Rockville, MD 20850, USA.

## Abstract

Human pluripotent stem cells (hPSCs) represent a powerful model system to study early developmental processes. However, lineage specification into trophectoderm (TE) remains poorly understood and access to well-characterized placental cells for biomedical research is limited, largely depending on fetal tissues or cancer cell lines. Here, we developed novel strategies enabling highly efficient TE specification that generates cytotrophoblast (CTB) and multinucleated primary syncytiotrophoblast (STB) followed by establishment of trophoblast stem cells (TSCs) capable of differentiating into extravillous trophoblast (EVT) and STB after long-term expansion. We confirmed stepwise induction of lineage- and cell-type-specific genes and substantiated typical features of placental cells using morphological, biochemical, integrated multi-omics, and single-cell analyses. Our data provide conclusive evidence that conventional hPSCs can be directly and exclusively converted into TE, thereby providing an unlimited source of diverse placental cell types suitable for a broad range of biomedical applications.

## INTRODUCTION

Inaccessibility of human tissue at early embryonic stages remains a major challenge for the study of placental development and function in health and disease. Typically, TSCs are isolated either from blastocysts or first-trimester placental villi^1, 2^. Derivation of TSCs from these sources is limiting for systematic studies and ethically problematic. Therefore, utilizing human induced pluripotent stem cells (iPSCs) as an inexhaustible source of self-renewing hTSCs and well-characterized placental cells would circumvent these issues, provide a tractable alternative to fetal tissues, and establish the ideal platform for translational studies.

A key event of early mammalian development is lineage segregation into TE and the inner cell mass (ICM). TE will generate the placenta as the critical organ of pregnancy, while the ICM gives rise to the epiblast (embryo proper) and primitive endoderm^3^. The current understanding is that *in vitro* culture of hPSCs generates cells with molecular features of primed pluripotency, resembling the post-implantation epiblast that can generate the primary embryonic germ layers (ectoderm, mesoderm, and endoderm) but not TE and trophoblast^3^. However, earlier studies suggested that primed or conventional hPSCs might differentiate into TE or express TE-associated genes upon treatment with BMP4 either alone^4–6^ or in combination with inhibitors of TGF-β and FGF signaling^7, 8^. Some reports suggested that these approaches may lead to only partial TE differentiation or induction of extraembryonic mesoderm^9–11^. More recent efforts to generate extraembryonic lineages from hPSCs focused on cells with enhanced potency, so-called naive pluripotency^12–15^ or isolation of TE-competent cells during cellular reprogramming into iPSCs^16, 17^.

Here, we report direct differentiation of hPSCs, routinely cultured in chemically defined E8 medium, into early TE cells and self-renewing TSCs that give rise to STB and EVT. Notably, our method exclusively generates cells of the TE lineage, whereas cell types of the embryonic germ layers are not produced. Moreover, the protocol is robust, reproducible with several hPSC lines, avoids genetic manipulation, and does not require the detour of inducing naive pluripotency. Taking advantage of this new approach, we provide deep cell characterization using integrated methods, perform multiple comparisons with other published datasets (including first-trimester placenta), and describe a comprehensive roadmap from pluripotency to defined placental cell types.

## RESULTS

### Exclusive lineage specification into TE

TE develops as an extraembryonic epithelial layer consisting of mononuclear CTB and multinucleated STB that anchor the implanting embryo to the decidua. To initiate TE differentiation, hPSCs cultured in E8 medium were switched to TE1 medium, which is E6 medium supplemented with a combination of A83-01, CHIR99021 (GSK-3 inhibitor), CH5183284 (FGFR inhibitor), BMP4, and BMP10 (**Fig. 1a).** The importance of using different BMP isoforms for the induction of trophoblast properties in hPSCs has been reported previously^4–7, 18^. TE1 culture conditions resulted in rapid morphological changes and emergence of cells with a cytoplasm appearing dark in phase-contrast brightfield images (**Fig. 1a**). Indeed, Periodic Acid-Schiff staining showed the cytoplasmic accumulation of large glycogen deposits, characteristic of early trophoblast^19^ (**Fig. 1b**). Withdrawal of BMPs, CH5183284, and an increased concentration of CHIR99021 on day 3 (switch to TE2 medium) then led to further epithelial differentiation and spontaneous formation of STB cells in a small subpopulation of cells (**Fig. 1a**). Subsequently, we established long-term, self-renewing TSC lines from the TE cells by switching to TE3 medium (described later in Figs. 3-5).

**Figure 1.**
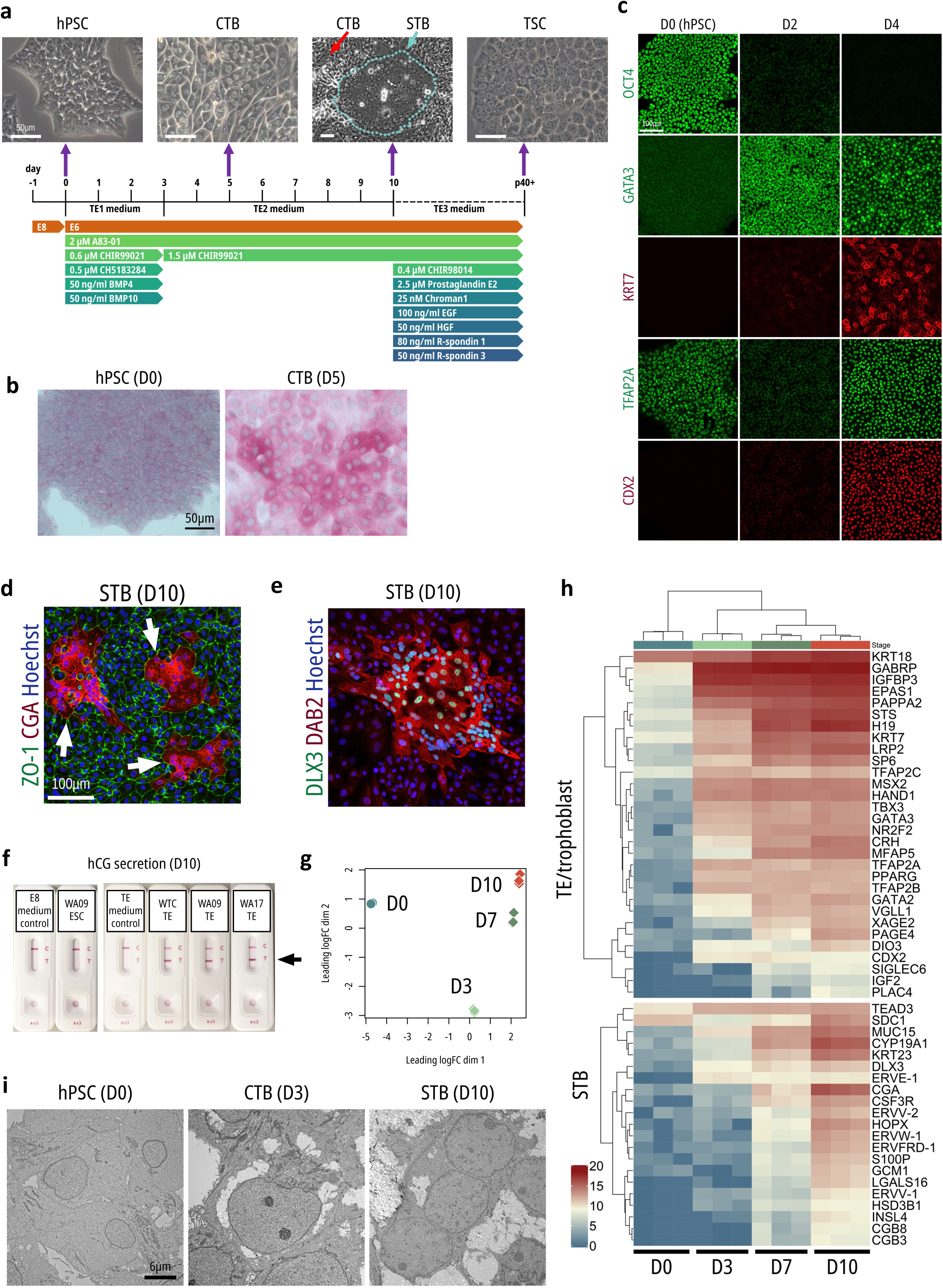
Morphology and characterization of early TE cells a, Overview of the roadmap that differentiated hPSCs into TE and self-renewing TSCs. Phase-contrast images correspond to select time points. **b**, Periodic Acid-Schiff (PAS) staining showing strongly increased glycogen storage in CTB versus undifferentiated cells. **c,** Immunocytochemistry showing direct differentiation of iPSCs (JHU198i) into TE. **d**,**e**, TE cultures at day 10 (D10) with marker expression characteristic of STB cells. **f**, hCG secretion into the culture medium was confirmed by a urine pregnancy test in D10 TE cells derived from 3 cell lines. **g**, MDS plot showing gradual changes in the transcriptional profiles of WA09 ESCs differentiating into TE. **h**, Heatmap of the selected most highly differentially expressed genes between D0 and D10 (log fold change > 8.5, FDR-adjusted P value < 1×10^-4^). **i**, Representative TEM images of WA09-TE cultures at D3 and D9 of differentiation along with ESC control (D0).

As early as day 2 after TE1 medium application, the pluripotency-associated transcription factor *OCT4* (*POU5F1*) was rapidly downregulated and *GATA3*, a pioneering TE specifier^4, 20, 21^, was strongly induced as shown by immunocytochemistry (**Fig. 1c**). Additional TE markers including *KRT7*, *TFAP2A*, and *CDX2*^4, 21^ were induced by day 4 (**Fig. 1c**). Between days 7-10, the monolayer of *ZO-1* expressing TE cells resembling CTB cells (**Fig. 1d**) spontaneously fused into large multinucleated STB-like structures (**Fig. 1a,d,e**; **Extended Data Fig. 1a-c, Supplementary Video)**, likely equivalent to the primary syncytium of the peri-implantation stage human embryo^22^. Human chorionic gonadotropin (hCG) is a critical pregnancy hormone and its α subunit, *CGA*, was specifically induced by STB (**Fig. 1d**). Similarly, STB markers *DAB2* and *DLX3*^23^ (www.proteinatlas.org) were detected by immunocytochemistry (**Fig. 1e**, **Extended Data Fig. 2b**). KRT18 is expressed in morula- and blastocyst-stage TE^24^ and was strongly expressed by both CTB and STB cells (**Extended Data Fig. 1c**). Next, the use of an over-the-counter pregnancy test confirmed secretion of hCG into the supernatants of TE cells derived from three hPSC lines at day 10 and its absence in human embryonic stem cell (hESC) cultures and medium controls lacking any cells (**Fig. 1f**). LC-MS/MS-based secretome analysis of supernatants confirmed enrichment of *CGB1* (chorionic gonadotropin subunit β1) and *CGA* peptide subunits in TE but not hPSC cultures (**Supplementary Table 1**).

Next, we performed time-course RNA sequencing (RNA-seq) analysis of hPSCs (D0) and differentiating TE cells harvested on days 3, 7, and 10. Multidimensional scaling (MDS) showed distinct clustering of samples across different timepoints (**Fig. 1h**). Early TE genes and STB markers were gradually upregulated (**Fig. 1g**, **Supplementary Table 2**), whereas pluripotency-associated genes were downregulated (**Extended Data Fig. 2a, Supplementary Table 2**). The pluripotency of undifferentiated cells lines was confirmed by PluriTest^25^ (www.pluritest.org) and as expected, TE samples were identified as non-pluripotent (**Extended Data Fig. 2b**).

Ultrastructural analysis using transmission electron microscopy (TEM) showed cell type-specific characteristics of pluripotent and differentiating cells. hPSCs displayed typical features of unspecialized cells, such as round, homogenous nuclei and few organelles surrounded by granular cytoplasm (**Fig. 1i**; **Extended Data Fig. 1d**). By day 3, CTB formed desmosomes, nuclei displayed prominent nucleoli, and their cytoplasm contained large amounts of rough ER (rER) and mitochondria. On day 10, large multinucleated STB structures were observed showing well-developed rER, mitochondria, microfilaments, Golgi, and large cytoplasmic vacuoles consistent with the identity and function of secretory placental cells.

Based on the conclusive results described above, we decided to revisit previous approaches, which used BMP4 alone or in combination with A83-01 and PD173074 (known as the BAP protocol) to induce trophoblast properties in hPSC^4–7^. When comparing these conditions to our TE1/TE2 method, all three treatments generated cells with distinct morphologies (**Extended Data Fig. 3a**) and widespread expression of *TFAP2A* (**Extended Data Fig. 3c**). Interestingly, TE1/TE2 media strongly induced expression of *CDX2*, which was absent in BAP-treated cultures and sporadically expressed in a few cells after application of BMP4 only (**Extended Data Fig. 3b**). Of note, BMP4 alone led to the induction of the mesodermal marker brachyury (*TBXT*) and showed residual expression of *OCT4*. Altogether, TE1/TE2 treatment was optimal for TE induction and our findings may in part explain the controversy in the literature when using BMP4 only or the BAP protocol.

### Transcriptional landscape of hPSC-derived TE cells

We performed detailed systematic RNA-seq experiments to analyze TE cells in comparison to the three primary germ layers derived from the same parental hESC lines (WA09, WA14, WA17) as well as two choriocarcinoma (CC) cell lines, JEG-3 and BeWo. MDS plots revealed that TE clustered away from hPSC and somatic lineages (**Fig. 2a**). CC lines clustered distinctly from all hPSC-derived cell types, likely reflecting their cancerous identities. Excluding CC lines from the MDS plot allowed visualization of the relationships between hPSC-derived cell types only, revealing a distinct path of the TE lineage (**Fig. 2b**). Among the most highly differentially expressed genes between hPSC and TE were typical trophoblast markers (**Supplementary Table 3**). STB markers were detected from the subpopulation of multinucleated STB cells (**Fig. 2c**). Gene set enrichment analysis of the TE-associated genes showed placenta and trophoblast as top terms based on three databases (**Fig. 2d**), while the most similar ARCHS4 cell lines identified were the CC lines BeWo and JEG-3 (**Fig 1e**). Downregulated genes in TE samples were markers of pluripotency, such as *OCT4*, *NANOG*, and *SOX2* (**Extended Data Fig. 4**). As expected, the three germ layers (ectoderm, mesoderm, endoderm) derived from hPSCs expressed typical markers. Genes such as *HAVCR1*, *DNMT3L*, and *GTSF1* were specifically expressed in the CC lines (**Extended Data Fig. 4**).

**Figure 2.**
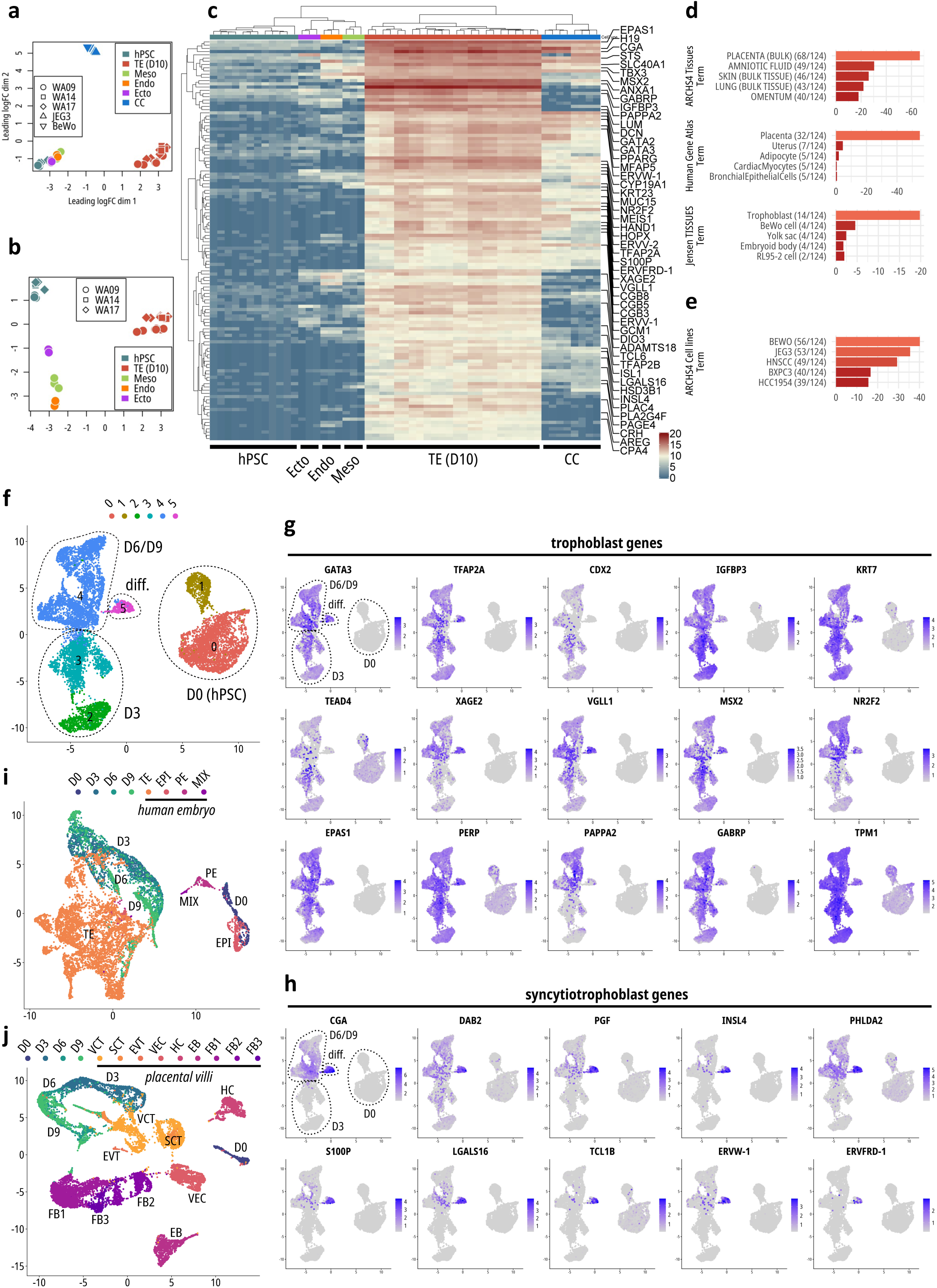
Analysis of the transcriptome of hPSC-derived TE cells. RNA-seq samples included TE derived from the three ESC lines WA09, WA14, and WA17 and D0 controls. WA09-derived endoderm (Endo), mesoderm (Meso), and ectoderm (Ecto) samples were included, as well as CC lines JEG-3 and BeWo. **a**,**b**, MDS plots of the above ESC lines **(a)** with and **(b)** without CC lines included. **c**, Heatmap of the 124 most highly differentially expressed genes (log-fold change > 8.5, FDR-adjusted P value < 1×10^-4^) between TE samples and hPSC samples. **d,** Enrichr analysis of the 124 differentially expressed genes displayed on the heatmap. **e**, Enrichr analysis – ARChS4 database. **f**, UMAP plot of time-course scRNA-seq of WA09 hPSC differentiation into TE. **g**, UMAP plots colored by the expression of individual differentially expressed TE or placenta-associated genes. **h**, Expression of markers associated with STB identity. **i**, Integration of the scRNA-seq data with data derived from human peri-implantation stage embryo^29^ using the integration method in R package “Seurat”. **j**, The same integration method with primary placental villi tissue^30^. VCT – villous cytotrophoblast, SCT – syncytiotrophoblast, EVT – extravillous trophoblast, FB – trophoblastic fibroblasts, VEC – vascular endothelial cells, EB – erythroblasts, HC – Hofbauer cells.

**Figure 3.**
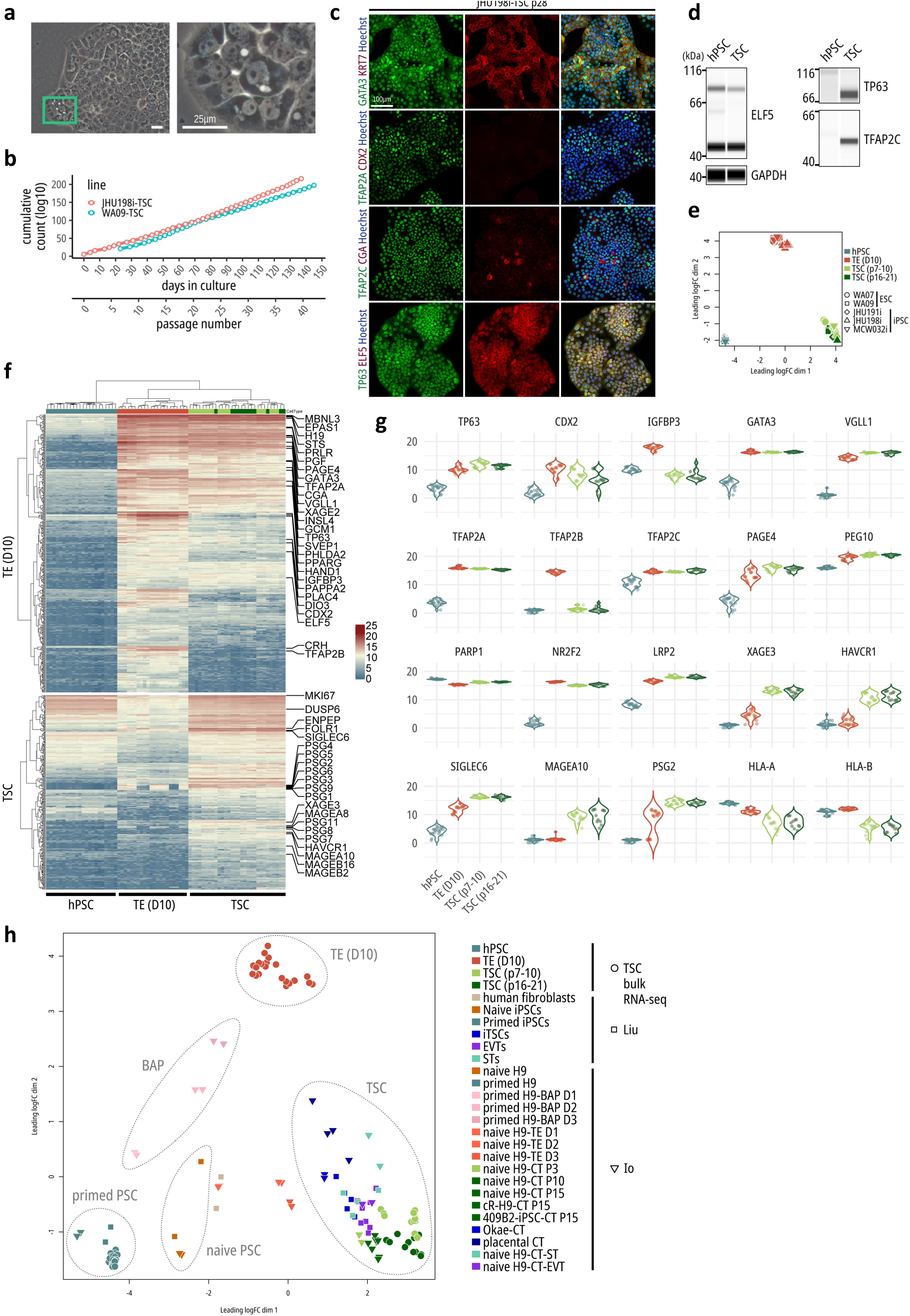
Establishment of self-renewing TSC lines from TE cells. **a**, Phase-contrast images of WA09-derived TSCs grown in TE3 medium. **b**, Passage-wise cumulative cell counts show stable proliferation of two TSC lines derived from one iPSC line (JHU198i) and one ESC line (WA09). **c**, Immunofluorescence showed the expression of TE/trophoblast markers in the JHU198i-TSC line at passage 28. **d**, Western blot analysis of TSC markers. **e**, RNA-seq MDS plot showed distinct clustering of hPSC, TE D10, and TSC samples, regardless of their genetic background. Early-passage (7-10) and late-passage samples were included (16-21). **f**, Heatmap of top differentially expressed genes (nearly 700) in TE D10 compared with hPSC (top, log2 fold change > 6, FDR-adjusted P value < 1×10^-4^) and in TSC compared with TE D10 (bottom, log2 fold change > 3, FDR-adjusted P value < 1×10^-4^). **g**, Expression of a number of markers associated with TE and trophoblast. **h**, Bulk RNA-seq integration with external datasets. MDS plot of transcriptional profiles of hPSC, TE (D10), and TSC and previously published iTSC^17^, naive hPSC-derived TSC, and placental TSC^15^. The original external sample group labels were maintained, however, the H9 ESC line is equivalent to WA09 presented here.

According to the Human Protein Atlas (HPA)^26–28,^ 91 genes and proteins are characteristic of the placenta. RNA-seq showed a high enrichment of these genes in hPSC-derived TE samples and to a lesser extent in CC lines while the number of these genes being expressed in hPSC and the three germ layers was low (**Extended Data Fig. 5**). Similarly, genes that were recently associated with TE of human peri-implantation stage embryos^29^ were enriched in hPSC-derived TE lines and CC lines but had limited expression in hPSC and the three germ layers (**Extended Data Fig. 6**).

### Single-cell analysis of TE derived from hPSCs

To characterize the differentiation process of pluripotent cells into TE, we performed time-course single-cell RNA-seq (scRNA-seq). The UMAP plot showed a clear separation between day 0 and day 3 samples, whereas day 6 and day 9 samples had more similar profiles (**Fig. 2f, Extended Data Fig. 7a**). Marker genes identified in each cluster (**Supplementary Table 4, Extended Data Fig. 7b**) included known pluripotency genes in day 0 cells (**Extended Data Fig. 7c**) and indicated acquisition of a TE signature in the remaining samples (**Fig. 2g**). Among the differentially expressed genes between hPSC (D0) and TE samples were early TE markers and HPA core 91 placenta-specific markers. Pan-trophoblast markers *KRT7* and *PERP*^30^ were strongly expressed. Cluster 5 (“diff.”) represented a subset of day 9 cells with markers of STB, including the two fusogenic syncytins *ERVW-1* and *ERVFRD-1*^31^ (**Fig. 2f, h**, **and Extended Data Fig. 7b**).

Next, integration of our dataset with a single-cell transcriptomic profile of human peri-implantation stage embryos^29^ revealed close clustering of the embryonic epiblast with the day 0 hPSC control and embryonic TE cells with day 3, 6, and 9 cells (**Fig 1i**). Another recent report focused on the first-trimester placenta and derived transcriptomic profiles of its single cells^30^. When compared to our dataset, day 0 (hPSC) samples clustered distinctly from all cell types and day 3, 6, and 9 cells were closest to the main primary trophoblastic cell types, which are villous cytotrophoblast (VCT), syncytiotrophoblast (SCT), and extravillous trophoblasts (EVT) (**Fig 1j**). These data suggest that hPSC-derived TE cells are most similar to the early TE of the human embryo rather than fully developed trophoblastic cell types of the placenta.

### Establishing self-renewing TSC lines

The CTB population in the early human placenta is thought to retain trophoblast stem or progenitor cell characteristics^2^. To establish TSC lines, we designed TE3 medium based on strong WNT pathway activation in combination with other factors and conditions that are important for placental development^1, 2, 27, 28, 32–34^ (**Fig. 1a**). We confirmed the strong expression of receptors for two key growth factors, EGF (*EGFR*) and HGF (*MET*), in TE cells at D10 (**Extended Data Fig. 8a**). WNT agonist CHIR99021 is toxic at higher concentrations^35^ and therefore we included the more potent GSK3-beta inhibitor CHIR98014^36^. Chroman 1 was recently reported as a more potent and specific ROCK inhibitor than Y-27632^37^ and was included in TE3 medium to prevent detachment of epithelial cells. Upon enzymatic dissociation of TE (day 10) and plating of small cell clumps into TE3 medium, colonies with proliferative cells formed and adopted a distinct morphology with prominent nucleoli and dark cytoplasm (**Fig. 1a, 3a**). Serial cell passaging was feasible and allowed the establishment of TSC cultures from two hESC and three hiPSC lines (**Fig. 3a,b and Extended Data Fig. 8b**). Self-renewing TSCs were capable of long-term expansion (> 5 months), remained undifferentiated, and expressed canonical trophoblast markers *GATA3*, *KRT7*, *TFAP2A*, *TFAP2C*, *TP63*, and *ELF5* (**Fig. 3c,d**). Interestingly, *CDX2* was either absent (**Fig. 3c**) or heterogeneously expressed (**Extended Data Fig. 9**) depending on the cell line, suggesting an important role for *CDX2* in early TE cells but not in TSCs. Western blot experiments revealed that *ELF5* was expressed in pluripotent cells and TSCs, whereas TP63 and TFAP2C were expressed in TSCs only (**Fig. 3d and Extended Data Fig. 8c**). Glycogen deposits observed in TE cells (**Fig. 1b**) were also detectable in TSCs (**Extended Data Fig. 8d**). Lastly, in contrast to hPSCs that were used as controls, established TSC lines did not pass the criteria for pluripotency as measured by PluriTest (**Extended Data Fig. 10a,b**).

### Analyzing the transcriptome and methylome of TSCs

We performed detailed RNA-seq experiments to characterize TSC lines and compare them to different developmental states. Parental hESC/iPSC lines, their differentiated progeny harvested as TE (D10), and established TSC lines showed distinct clustering in the PCA plot (**Fig. 3e**). The transition from pluripotency to TE (D10) resulted in extensive global transcriptional changes (**Fig. 3f**, **Supplementary Table 5**), reflecting our earlier analysis (**Fig. 2a-e**). Further changes were observed when TE (D10) was differentiated in TSCs (**Fig. 3f**). Multiple trophoblast markers were upregulated and expressed in a cell-type-specific fashion (**Fig. 3g**). Selective markers of primary first-trimester VCT cells (*PAGE4*, *PEG10*, and *PARP1*)^30^ and markers associated with stemness of human TSC (*NR2F2*, *LRP2*, and *PEG10*)^16^ were upregulated or highly expressed in TSC. The top 3 genes with the highest increase in expression in TSC over TE (D10) were *XAGE3*, *HAVCR1*, and *MAGEA10*. Pregnancy-specific glycoproteins were upregulated in TSC after an incomplete upregulation in TE (D10) samples. HLA-A and -B were downregulated in TSC compared with TE (D10) and hPSC control (**Fig. 3g**). The lack of HLA-B expression in TSC was confirmed by qRT-PCR in comparison with human dermal fibroblasts and was comparably low in the JEG-3 cell line (**Extended Data Fig. 11c**). This downregulation represents one of the molecular trophoblast criteria^11^. HAVCR1 was recently implicated as a marker of TE and was absent in amnion, while IGFBP3 expression followed the opposite trend^14^. *IGFBP3* was strongly upregulated in TE (D10) and returned to its basal level observed in hPSC upon establishment of TSC. *SIGLEC6*, a marker of CTB cells in the human chorionic villi at week 5, cultured human placenta-derived TSC, and naive human PSC-derived TSC^15^, was moderate in TE (D10) but high in TSC (**Fig. 3f,g**).

A direct comparison and integration of RNA-seq data with previously published datasets^15, 17^ showed that our hPSC samples (D0) clustered closely with primed hPSCs and distinctly from cells with naïve pluripotency. The TSCs derived with our method clustered closely with all trophoblastic cell types, including TSCs derived from first-trimester placental villi. Primed hPSC treated with the BAP protocol did not cluster with any other samples supporting the notion of incomplete differentiation. Interestingly, regardless of whether TSCs were derived from primed or naive hPSCs, they converged on a phenotype with very similar transcriptomic profiles (**Fig. 3h**). In other words, the TE lineage can be directly generated from routinely cultured primed hPSCs without the induction of naïve pluripotency.

Next, scRNA-seq of TSCs (WA09) identified 4 cell clusters (**Fig. 4a**). The majority of cells expressed *PAGE4*, *PEG10*, *PARP1*, *NR2F2*, and *LRP2*. Trophoblast marker VGLL1^16^ was expressed at lower levels in cluster 1, which had an elevated expression of *VIM* and *COL1A2*. *CGA*, *INSL4*, and other STB marker expression was characteristic of cluster 3 and represented a small subpopulation of cells with a spontaneous propensity towards fusion (**Fig. 4b**). Genes identified as markers of each cluster are summarized in **Supplementary Table 6**.

**Figure 4.**
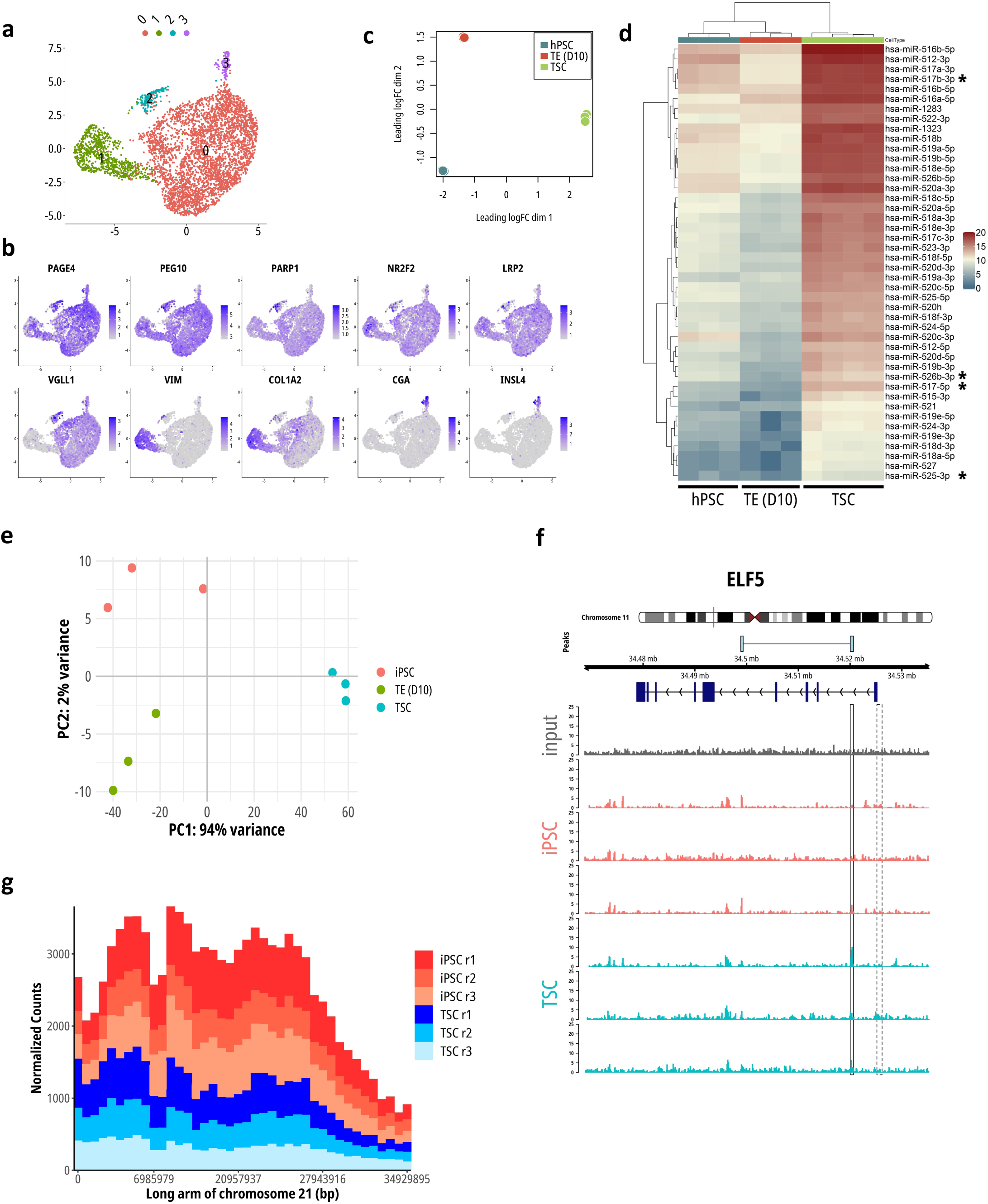
Single cell transcriptomics, miRNA expression, and epigenetic analysis of TSCs confirming trophoblast criteria. **a**,**b,** Single cell analysis (scRNA-seq) of TSCs (WA09) identified 4 main clusters. Trophoblast markers expressed by CTB of the placental villi and STB. **c**, MDS plot of miRNA expression profiles show distinct clustering of hPSC, TE D10, and TSC (JHU198i line). **d**, Heatmap of the expression of microRNAs of the C19MC cluster. The miRNAs marked by asterisks confirm key trophoblast criteria according to ref. 10. **e**, Principal component analysis of methylation profiles in iPSCs (JHU198i), TE (D10), and TSC based on MeDIP-seq. **f**, Absence of methylation in the ELF5 promoter region (dashed line) was found across samples. **g**, Low methylation levels along the long arm of chromosome 21 (q21) are consistent with trophoblast identity according to refs. 2 and 37.

**Figure 5.**
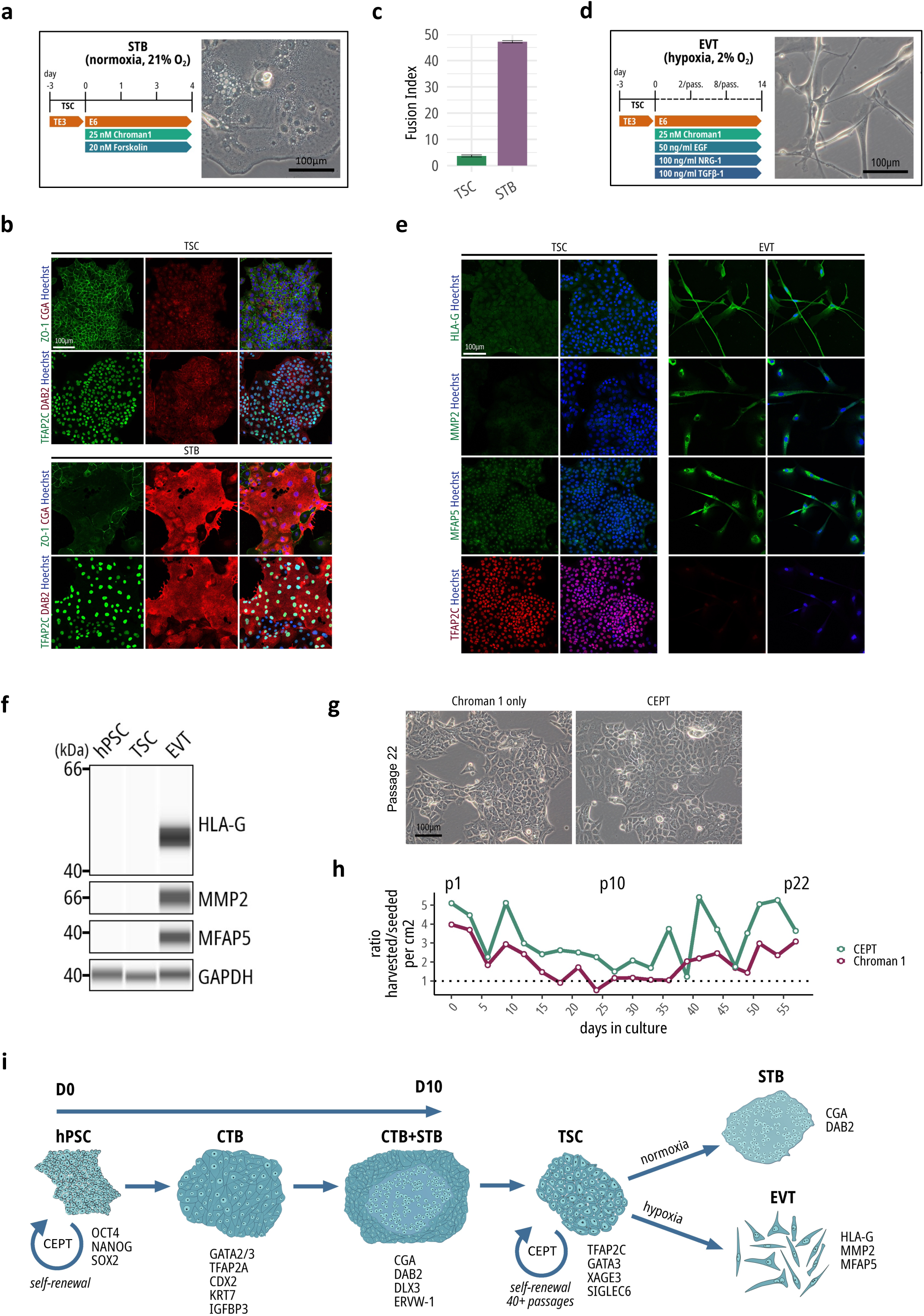
Terminal differentiation of TSCs and enhancement of TSC propagation by using CEPT. **a**, Protocol for differentiation of TSC into STB over roughly 4 days, leading to the formation of multinucleated cells. **b**, Immunofluorescence of STB markers *CGA* and *DAB2*, TSC marker *TFAP2C*, and gap junction protein *ZO-1* (*GJP1*) delineating individual cellular borders. **c**, Fusion index, which is the proportion of nuclei in multinucleated STB cells relative to all nuclei, was calculated from the fluorescent images. **d**, Protocol for differentiation of TSC into EVT over 14 days with passaging (“pass.”) on days 2 and 8, and typical cell morphology. **e**, Immunofluorescence analysis of EVT markers in WA09-TSC-derived EVT. **f**, Western blot analysis of EVT marker expression in hPSC, TSC, and EVT (WA09). **g**, Representative phase-contrast images of TSC lines derived using CEPT compared to control (Chroman 1 only). **h,** Ratio of the number of harvested cells per cm^2^ divided by the number of seeded cells per cm^2^ at each passage, in CEPT-treated and control (Chroman 1 only) TSC. **i,** Summary of placental cell development recapitulated by directed differentiation of hPSCs into distinct cell types expressing specific markers and showing distinct morphologies.

Micro-RNA sequencing (miRNA-seq) revealed distinct clustering of hPSC, TE (D10), and TSC (**Fig. 4c**, **Extended Data Fig. 12a**) and upregulation of all C19MC miRNAs in TSCs, which is a critical trophoblast criterion^11^, compared with hPSC and TE (D10) controls (**Fig. 4d**, **Extended Data Fig. 12b**). Similarly, because of its importance as a defining trophoblast feature, we analyzed *ELF5* methylation status as well as the global methylome of the hPSC-derived TSCs by using methylated DNA immunoprecipitation sequencing (MeDIP-seq). Undifferentiated iPSCs and TE (D10) showed smaller differences, but the cells underwent dramatic changes in DNA methylation following the establishment of TSCs (**Fig. 4e, Extended Data Fig. 13a**). We, therefore, focused on the comparison of pluripotent cells versus TSCs. The *ELF5* promoter was unmethylated in TSC, a trophoblast criterium^11^ (**Fig. 4f, Extended Data Fig. 13b**). Notably, unlike a previous report^11^, the ELF5 promoter was unmethylated in hPSC, which may confer TE competency to primed human cells identified in the present study. On the protein level, *ELF5* was expressed in both hPSC and TSC (**Fig. 3c, d**; **Extended Data Fig. 8c**). On the transcript level, *ELF5* expression in TE (D10) was comparable with CC cell lines (**Extended Data Fig. 11a**) and was maintained in TSC (**Extended Data Fig. 11b**) at a level comparable with the JEG-3 cell line **(Extended Data Fig. 11c**).

TSCs were previously reported to have lower levels of methylation along the long arm of chromosome 21^2, 38^. We confirmed this feature in hPSC-derived TSCs (**Fig. 4g, Extended Data Fig. 13c**). The MeDIP-seq global methylation changes were summarized for further analysis in the form of heatmaps (**Extended Data Fig. 14a,b**) and Manhattan plots (**Extended Data Fig. 14c,d**) of the most highly differentially enriched peaks (**Supplementary Table 7**); and locus plots of the top 8 differentially methylated genes (**Extended Data Fig. 15a,b**).

### Differentiation of TSCs into terminal cell types

To differentiate TSC into STB, we applied a simple culture condition that includes forskolin^2^ and Chroman 1 (**Fig. 5a**). Over the course of differentiation, multinucleated cells formed and expressed STB markers CGA and DAB2 (**Fig 4b**, **Extended Data Fig. 16, 17**). ZO-1 expression disappeared between the individual cells upon fusion, whereas TFAP2C remained expressed in both TSC and STB. After 4 days in culture, almost 50% of the nuclei were embedded in multinucleated cells (**Fig. 5c**).

We attempted to apply a previously published method for differentiation of TSCs into EVT^2^ that included NRG1, however, we were unable to reproduce the epithelial-to-mesenchymal transition (EMT) and formation of cells with spindle-shaped morphology. To induce efficient EMT, we designed a novel EVT medium containing a combination of EGF, NRG1, TGFβ1, and Chroman 1 (**Fig. 5d**). TGFβ1 was included because it is a well-known inducer of EMT^39^, although there are conflicting reports on its role in EVT differentiation^40^. In addition to the EVT medium composition, incubation in hypoxic (2%) culture was used as it was previously reported to enhance differentiation of TSC into EVT^41^. When exposed to these conditions, TSCs underwent EMT and acquired spindle-shaped morphologies (**Fig. 5d**). Immunofluorescence imaging (**Fig. 5e**, **Extended Data Fig. 18**) and Western blot analysis (**Fig. 5f**) confirmed expression of the EVT markers in these cells and their absence in hPSC and TSC controls. Hence, functional differentiation of TSC into terminal cell types with distinct molecular and cellular features provided additional strong evidence for their biological identity.

Next, to optimize cell viability during TSC line expansion, we incorporated the recently developed small-molecule cocktail “CEPT”^37^, which has been shown to have beneficial effects on cell survival and cytoprotection of hPSCs and multiple other cell types. CEPT treatment for 24 h at each passage led to a significant reduction in the proportion of dead cells as measured over multiple passages and compared to the application of the ROCK pathway inhibitor Chroman 1 only (**Fig. 5g,h, Extended Data Fig. 19a,b**). These observations were replicated in an additional instance of TSC establishment from WA09 (**Extended Data Fig. 20a**) and two iPSC lines (**Extended Data Fig. 20b,c**). To summarize all results, we propose a defined *in vitro* roadmap for the human trophectoderm lineage that recapitulates hallmarks of human placental development (**Fig. 5i**).

## DISCUSSION

Here, we demonstrate that conventional hESC and iPSC lines can be directly, reproducibly, and exclusively differentiated into the TE lineage producing distinct functional phenotypes including CTB, STB, EVT, and self-renewing TSCs. Our findings provide a new perspective as the current understanding is that only naive pluripotency, thought to correspond to pre-implantation epiblast, retain the capacity for TE differentiation. Accordingly, only naive hPSCs could be differentiated into self-renewing TSC when using specific culture conditions^2^, whereas primed cells only generated non-self-renewing CTB in response to BMP4 but failed to generate TSCs^13^. Interestingly, another study suggested that primed hPSCs can differentiate into amniotic epithelium but not TE^14^. These observations were replicated in a concurrent study and the resulting TSCs passed the trophoblast criteria, whereas primed hPSCs incompletely differentiated into TE-like cells using the BAP protocol did not^15^. A more complex approach for deriving human TSCs is to isolate a subpopulation of cells with trophoblast properties that emerged during the reprogramming of fibroblasts into iPSCs^16, 17^. Considering our present findings, it is likely that previous attempts were insufficient for generating *bona fide* trophoblast from primed hPSCs. The first report describing the possibility of TE competency of hESCs by Thomson and colleagues^5^ spurred two decades of controversy. The robustness of our methods, firmly based on using chemically defined conditions, provides conclusive evidence that routinely cultured hPSCs can indeed be directly coaxed into TE, while other developmental lineages were not induced. We envisage that the detailed characterization experiments and rich datasets that we established here will serve as an invaluable resource for studying lineage commitment of embryonic and extra-embryonic tissues. Most importantly, our study provides the framework for utilizing human placental cells from an inexhaustible source such as iPSCs for pioneering translational experiments to better understand and treat infertility and gestational diseases.

## Supporting information

Ext Data Fig 1

Ext Data Fig 2

Ext Data Fig 3

Ext Data Fig 4

Ext Data Fig 5

Ext Data Fig 6

Ext Data Fig 7

Ext Data Fig 8

Ext Data Fig 9

Ext Data Fig 10

Ext Data Fig 11

Ext Data Fig 12

Ext Data Fig 13

Ext Data Fig 14

Ext Data Fig 15

Ext Data Fig 16

Ext Data Fig 17

Ext Data Fig 18

Ext Data Fig 19

Ext Data Fig 20

## Acknowledgements

We thank all our colleagues at NCATS for their contributions. This work utilized the computational resources of the NIH HPC Biowulf cluster (http://hpc.nih.gov). MeDIP-seq analysis and single-cell data integration was completed in part by Rancho Biosciences. Graphic illustrations were designed by the NIH Medical Arts Branch. The authors also thank Hannah Baskir for editing the manuscript and suggestions.

## Author contributions

J.S. and I.S. conceived the project. Experiments: J.S., C.A.T.,S.R., P-H.C., T.D., Y.G., P.O. Data analysis and discussions: J.S., C.A.T., S.R., P-H.C., T.D., Y.G., P.O., S.M., T.C.V., A.S., I.S. Manuscript writing: J.S. and I.S.

**Extended Data Figure 1. Fusion of CTB into primary STB and ultrastructural analysis of differentiating cells**

**a**, Additional phase-contrast images of STB cells of different sizes after 10 days of TE differentiation. **b**, Example of *DLX3* immunofluorescence exclusive to the nuclei of STB cells, overlaid on a phase-contrast image. **c**, *KRT18* immunofluorescence, observed in all TE cells, including STB cells. **d**, Related to Figure 1h. The images show gradual change in ultrastructure from undifferentiated, stem cell-like properties in hPSC to more defined and mature organelles at D10.

**Extended Data Figure 2. Expression of pluripotency-associated genes during early TE differentiation**

**a**, Heatmap of 138 of the most highly differentially downregulated genes (log fold change < -7.5, FDR-adjusted P value < 1×10^-4^) between D0 and D10 of differentiation in the WA09 line. **b**, PluriTest analysis of the global transcriptional profiles of D0 hPSC and D10 TE samples.

**Extended Data Figure 3. Comparison of TE induction method to BMP4 treatment and BAP protocol.**

**a**, Representative phase-contrast images. **b**, Imaging based on immunocytochemistry. TE1/TE2 – TE induction protocol presented in this study, “BAP” protocol – treatment with E6 basal medium + BMP4 + A83-01 + PD173074.

Extended Data Figure 4. Bulk RNA-seq heatmap of differentially expressed genes in cell types other than TE.

Related to Figure 2 bulk RNA-seq, which was the source of the data. Roughly 20-40 most highly differentially expressed genes representing markers of each cell type were included. The FDR-adjusted P value for all genes was less than 1×10^-4^. The log fold change cut-off for hPSC was 8, Ecto 7, Endo 9, Meso 9, CC 8. “hPSC” – human pluripotent stem cells, “Ecto” – hPSC-derived ectoderm, “Endo” – hPSC-derived endoderm, “Meso” – hPSC-derived mesoderm, “TE D10” – hPSC-derived TE at day 10, “CC” – choriocarcinoma cell lines JEG-3 and BeWo.

**Extended Data Figure 5. Bulk RNA-seq heatmap of Protein Atlas placental gene expression.**

Related to Figure 2 bulk RNA-seq, which was the source of the data. Protein Atlas placenta-enriched genes (91 genes) were included. “hPSC” – human pluripotent stem cells, “Ecto” – hPSC-derived ectoderm, “Endo” – hPSC-derived endoderm, “Meso” – hPSC-derived mesoderm, “TE D10” – hPSC-derived TE at day 10, “CC” – choriocarcinoma cell lines JEG-3 and BeWo.

**Extended Data Figure 6. Bulk RNA-seq heatmap of the expression of previously identified human TE genes.**

Related to Figure 2 bulk RNA-seq, which was the source of the data. TE-specific genes^29^ were included. “hPSC” – human pluripotent stem cells, “Ecto” – hPSC-derived ectoderm, “Endo” – hPSC-derived endoderm, “Meso” – hPSC-derived mesoderm, “TE D10” – hPSC-derived TE at day 10, “CC” – choriocarcinoma cell lines JEG-3 and BeWo.

**Extended Data Figure 7. Single cell analysis of hPSC-derived TE cells.**

Additional plots related to Figure 2 scRNA-seq. **a**, UMAP plot colored by the original samples. **b**, Heatmap of top 10 markers of individual clusters. **c**, Expression of pluripotency markers, mostly confined to clusters 0 and 1, identified as D0 hPSCs.

**Extended Data Figure 8. Additional data confirming TSC establishment.**

**a**, Expression of receptors for growth factors EGF (*EGFR*) and HGF (*MET*), based on RNA-seq (Fig. 2 bulk RNA-seq dataset). **b**, Phase-contrast images of proliferating TSC lines derived from three hiPSC lines (JHU191i, JHU198i, MCW032i) and an additional hESC line (WA07) with similar morphological features. **c**, Related to Fig. 3c. Control hPSC cultures were immunostained for TSC markers. **d**, Periodic Acid-Schiff staining of TSC colonies showing a representative example. The pink color marks the glycogen deposits in the cytoplasm of cells.

**Extended Data Figure 9. Immunocytochemical analysis of TSC line derived from hESCs (WA09).**

The presented data is related to Fig. 3c. TSC line was derived from WA09 cells and analyzed at passage 12.

**Extended Data Figure 10. Expression of pluripotency-associated genes in hPSC-derived TSCs.**

**a**, Immunocytochemical analysis of pluripotency markers *OCT4* and *NANOG* in pluripotent cells and TSCs derived from an iPSC line (JHU198i). **b**, PluriTest analysis of the global transcriptional profiles of TSC lines and D0 hPSC controls.

**Extended Data Figure 11. Expression of ELF5 in hPSC-derived TE and TSCs.**

**a**,**b**, Levels of *ELF5* in samples of Fig. 2 bulk RNA-seq (**a**) and Fig. 3 bulk RNA-seq (**b**), compared with *GATA3*. **c**, qRT-PCR in TSC markers and *HLA-B* in fibroblasts, WA09-derived TSC and JEG-3 CC line. y-axis values were reversed delta CT values (40 – delta CT) that represent CT values normalized to *GAPDH*. Two biological and 4 technical replicates were used. Significance tests obtained by linear modeling were summarized in the table below. “adj. P value” – adjusted P value.

**Extended Data Figure 12. miRNA-seq analysis of an additional TSC line.**

**a**, MDS plot of hPSC, TE D10, and TSC (JHU191i line) based on their profiles of miRNA expression. **b**, Heatmap of the expression of miRNAs of the C19MC cluster. The miRNAs marked by asterisks were previously preferentially studied as part of defining the trophoblast criteria.

**Extended Data Figure 13. MeDIP-seq analysis in additional TSC line.**

**a**, Principal component analysis of methylation profiles of JHU191i hPSC, TE (D10), and TSC based on MeDIP-seq. **b**, No evidence of methylation in the *ELF5* promoter region (dashed line) was found across samples. **c**, Levels of methylation along the long arm of chromosome 21 (q21).

**Extended Data Figure 14. MeDIP-seq differential enrichment analysis in TSC**

**a**,**b**, Heatmaps of peaks identified as differentially enriched between iPSC and TSC (lines JHU198i and JHU191i). The peaks were associated with the genes displayed on the heatmap. The colors correspond to the level of methylation, increasing with the values on the color key. **c**,**d**, Manhattan plots of identified peaks differentially enriched between iPSC and TSC. The most significant peaks were highlighted and colored based on their methylation either in iPSC or TSC.

**Extended Data Figure 15. MeDIP-seq locus plots of identified peaks differentially enriched between iPSC and TSC.**

**a**,**b**, Presented genes were the most highly differentially methylated between iPSC and TSC generated from JHU198i (**a**) and JHU191i (**b**) iPSC lines. The dashed lines denote the promoter regions.

**Extended Data Figure 16. Immunofluorescence analysis of ZO-1 and CGA in TSC-derived STB.** Additional examples related to Fig. 5b.

**Extended Data Figure 17. Immunofluorescence analysis of TFAP2C and DAB2 in TSC-derived STB.** Additional examples related to Fig. 5b.

**Extended Data Figure 18. Immunofluorescence analysis of HLA-G in TSC-derived EVT.** Additional examples related to Fig. 5e.

E**xtended Data Figure 19. Enhancement of TSC viability with CEPT.**

**a**, The percentage of dead cells in WA09-TSC (stained with propidium iodide, red fluorescence) out of all cells in Chroman 1 and CEPT-treated TSCs. Viable cells were stained with Calcein AM (green fluorescence). **b**, Representative Incucyte fluorescent images of viable and dead cells in Chroman 1- and CEPT-treated TSC.

**Extended Data Figure 20. Immunofluorescence analysis of TSC marker expression in additional lines of CEPT-treated TSC.**

**a**-**c**, Additional TSC lines were established using Chroman 1 alone (control) and CEPT treatment during the first 24 h following passaging from WA09 ESC (**a**), NL5 iPSC (**b**), and LiPSC-GR1.1 iPSC (**c**) at passage 13, 9, and 7, respectively.

Supplementary Video. Time-lapse phase-contrast imaging of iPSCs (JHU198i) differentiating into TE.

The cells were cultured in TE1 medium until day 3, then switched to TE2 medium. Imaging was started on day 4 and continued for 9 days. Images were taken every 4 h. A subset of mononucleated cells fused into multinucleated STB cells. The experiment was done using an Incucyte instrument.

**Supplementary Table 1.** Nano LC-MS/MS secretome analysis of hPSC-derived trophectodermal cells. The secretome of hPSC-derived TE cells at D10 was compared with hPSC secretome.

**Supplementary Table 2.** Results of the differential expression analysis of RNA-seq data presented on Fig. 1.

**Supplementary Table 3.** Results of the differential expression analysis of bulk RNA-seq data presented on Fig. 2.

**Supplementary Table 4.** Marker genes of each identified Fig. 2 scRNA-seq cluster, ordered for each cluster based on evidence of differential gene expression (P value).

**Supplementary Table 5.** Results of the differential expression analysis of bulk RNA-seq data presented on Fig. 3.

**Supplementary Table 6.** Marker genes of each identified Fig. 4 scRNA-seq cluster, ordered for each cluster based on evidence of differential gene expression (P value).

**Supplementary Table 7.** Results of the differential enrichment analysis for MeDIP-seq data, with differentially methylated regions annotated based on overlaps with known genes or non-coding sequences.

**Supplementary Table 8.** List of reagents and antibodies used in this study.

**Supplementary Table 9.** scRNA-seq sequencing parameters for Figs. 2 and 4.

## METHODS

### hPSC culture

All hESCs (WA07, WA09, WA14, and WA17; WiCell) and hiPSCs (JHU191i, JHU198i, and MCW032i; WiCell; WTC iPSC; Allen Institute for Cell Science) were maintained under feeder- and xeno-free conditions in Essential 8 (E8) medium (A1517001, Thermo Fisher Scientific) on microplates or T175 flasks coated with vitronectin (VTN; A14700, Thermo Fisher Scientific) at a concentration of 0.5 µg/cm^2^. Cells were passaged every three days. The hPSC colonies were treated with 0.5 mM EDTA (15575020, Invitrogen) in phosphate buffered saline (PBS) without calcium or magnesium (14190144, Gibco) for 5-6 min to dissociate the hPSC colonies. The resulting cell clumps were counted using the Nexcelom Cellometer automated cell counter. The clumps were then plated at a density of 2-3 × 10^5^ cells per cm^2^ in E8 medium and maintained in a humidified atmosphere containing 5% CO_2_ at 37°C.

Where indicated, hPSC were cultured in presence of a cocktail of compounds 50 nM Chroman 1 (HY-15392, MedChemExpress), 5 µM Emricasan (S7775, Selleck Chemicals), 1× Polyamine Supplement (P8483-5ML, Millipore-Sigma), and 0.7 µM trans-ISRIB (5284, Tocris Biosciences) termed “CEPT”, up to 24 h following passaging to enhance the viability of the cells. In the case of TSC culture, TE3 medium contains 25 nM Chroman 1 and this concentration is increased to 50 nM at each passage, with the remaining “EPT” components added for a full cocktail.

### Differentiation of hPSCs into TE and proliferative TSCs

hPSC maintained in E8 medium on VTN-coated 6-well plates, as described above, were dissociated using 0.5 mM EDTA for 5-6 min. The resulting clumps were then counted using the Nexcelom Cellometer and seeded at a density of 4-5 × 10^5^ per cm^2^ in E8 medium on VTN-coated 6-well plates. 24 h later (day 0), the spent medium was replaced with TE1 medium and changed every day until day 3. On day 3, the medium was replaced with TE2 medium until day 7-10, with daily medium changes. Then the monolayers containing a mixture of mononucleated CTB-like cells and multinucleated STB cells were partially dissociated using StemPro Accutase (A1110501, Gibco) for 8-12 minutes. The resulting clumps of cells were plated at a density of 2.5 × 10^5^/cm^2^ in TE3 medium supplemented with additional 25 nM of Chroman 1. 24 h later, the medium was changed with fresh TE3 medium with basal concentration of Chroman 1 (25 nM). The subsequent media changes were performed daily. The subsequent passaging steps were performed using Accutase treatment for 5-8 min. The seeding density remained at 2.5 × 10^5^/cm^2^ for an additional passage. Then the seeding density was increased to 3-4 × 10^5^/cm^2^ and remained the same for all passages on. The passaging interval was 3 days and the derived proliferative cell lines were passaged over 40 times. Cells were maintained in a humidified atmosphere containing 5% CO_2_ at 37°C.

Where indicated, for the first 24 h of culture following passaging, the use of 50 nM of Chroman 1 was complemented with 5 μM Emricasan, polyamines, and 0.7 μM trans-ISRIB to make up the full CEPT cocktail for viability enhancement, as described above for the culture of hPSCs.

### Time-lapse phase-contrast imaging of cellular fusion

hPSC were differentiated into TE on VTN-coated Incucyte ImageLock 96-well plates (4379, Sartorius) according to the TE differentiation protocol described above. Phase-contrast imaging was performed using the Incucyte S3 (Sartorius) starting on day 4 with images taken every 4 h over the course of 9 days.

### Differentiation of TSCs into EVT and STB

For EVT differentiation, TSC were cultured in TE3 medium in hypoxia (2% O_2_) for at least 1 passage (3 days). Then the cultures were dissociated with Accutase for 10-15 min and seeded at a density of 6 × 10^5^/cm^2^ in EVT medium on VTN-coated culture plates. After 2 days of culture, the cells were dissociated with Accutase, replated at a density of 2 × 10^5^/cm^2^ in EVT medium, and cultured for another 6 days. Some cell lines required additional passages with 6 days intervals. The EVT medium was changed every other day and the cells were incubated at 37°C, 5% CO_2_, and 2% O_2_.

For STB differentiation, TSC cultures were dissociated with Accutase for 5-8 min, seeded at a density of 4-5 × 10^5^/cm^2^ in STB medium, and cultured for 3-4 days at 37°C, 5% CO_2_, and 21% O_2_.

### Fusion index of TSCs differentiated into STB

Fluorescent images were taken on the high-content imaging platform Opera Phenix (PerkinElmer). Image analysis was performed using the online interface of the Columbus software (PerkinElmer). *ZO-1* (*TJP1*) expression was used to guide the categorization of cells into mononucleated and multinucleated. CGA expression was used to guide the identification of multinucleated cells. A total of 225 fields across two wells were analyzed. Fusion index was calculated as the percentage of nuclei found in multinucleated cells out of all nuclei.

### Culture of human dermal fibroblasts

Human dermal fibroblasts (PCS-201-012, ATCC) were seeded on tissue culture-treated 6-well plates at a density of 5,000 cells/cm^2^ in fibroblast basal medium (PCS-201-030, ATCC) supplemented with components of the Fibroblast Growth Kit, Low Serum (PCS-201-041, ATCC) and incubated in a humidified atmosphere at 37°C, 5% CO2, and 21% O2. Medium was changed daily, and the cells were passaged every three days by treatment with Accutase for 6 min, followed by dilution with culture medium, centrifugation at 200 g for 3 min, and replating onto 6-well plates.

### Culture of choriocarcinoma cell lines

Human CC cell lines JEG-3 (HTB-36, ATCC) and BeWo (CCL-98, ATCC) were seeded on tissue culture-treated 6-well plates at a density of 20,000 cells/cm^2^ in Eagle’s Minimum Essential Medium (30-2003, ATCC) and Kaighn’s Modification of Ham’s F-12 Medium (30-2004, ATCC), respectively. Both JEG-3 and BeWo cell lines were supplemented with 10% fetal bovine serum (FBS; 30-2020, ATCC). The cultures were incubated in a humidified atmosphere at 37°C, 5% CO_2_, and 21% O_2_. Medium was changed daily and the cells were passaged every three days by treatment with Accutase for 15 min, followed by dilution with culture medium, centrifugation at 200 g for 3 min, and replating onto 6-well plates.

### Differentiation of hPSC into somatic lineages

Differentiation of hPSC into endoderm was initiated using the STEMdiff™ Definitive Endoderm Kit, TeSR™-E8™ Optimized (05115, STEMCELL Technologies). Cells were plated at a density of 150,000 cells/cm^2^ on 6-well plates in E8 media supplemented with 50 nM of the ROCK inhibitor Chroman 1. After reaching 50-60% confluency, cell culture media was switched to TeSR-E8 Pre-Differentiation media for 24 h at which the cells reached 70% confluency. After aspirating the culture medium, cells were dissociated into single cells by 10-15 min incubation with 0.5 mM EDTA (15575020, Thermo Fisher Scientific) in PBS without calcium or magnesium (14190144, Thermo Fisher Scientific) at 37°C and plated at a density of 210,000 cells/cm^2^ onto VTN-coated 6-well plates in TeSR-E8 Pre-Differentiation media supplemented with 50 nM Chroman 1. 24 h post plating, the cultures were rinsed with DMEM/F-12 (10565018, Gibco) and media was replaced with Medium 1 (STEMdiff Definitive Endoderm Basal Medium with STEMdiff Definitive Endoderm Supplement A and STEMdiff Definitive Endoderm Supplement B). The next day, cell culture media was exchanged with Medium 2 (STEMdiff Definitive Endoderm Basal Medium with STEMdiff Definitive Endoderm Supplement B). On days 3-5, cell culture media was changed daily with Medium 2 (STEMdiff Definitive Endoderm Basal Medium with STEMdiff Definitive Endoderm Supplement B). On day 5, cells were ready for end-point assay.

Mesoderm differentiation of hPSC was induced using the STEMdiff Mesoderm Induction Medium (05221, STEMCELL Technologies). Cells were plated at a density of 50,000 cells/cm^2^ on VTN-coated 6-well plates in E8 media supplemented with 50 nM Chroman 1 and incubated for 24 h. On days 2-5, cell culture medium was replaced with STEMdiff Mesoderm Induction Medium. On day 5, cells were ready for end-point assay.

For ectoderm differentiation, hPSC were plated at a density of 50,000 cells/cm^2^ on VTN-coated 6-well plates in E8 media supplemented with 50 nM Chroman 1. 24 h later, the cell culture media was switched to E6 (A1516401, Thermo Fisher Scientfic) supplemented with 100 nM LDN 193189 dihydrochloride (6053, Tocris) and 2 µM A83-01. Media was changed daily for 6 days. On day 7, cells were ready for end-point assay. For all differentiation protocols, cells were maintained at 37 °C in a humidified atmosphere containing 5% CO_2_ and 21% O_2_.

### Cryopreservation

Both hPSC and TE cells were harvested using EDTA and Accutase, respectively, as described above. The cell suspensions were centrifuged at 200 g for 3 min and resuspended in CryoStor CS10 (210102, BioLife Solutions) and placed into -80°C freezer in CoolCell containers (432000, Corning) overnight, then placed into -150°C freezer for long-term storage.

### Immunocytochemistry

hPSC, hPSC-derived TE cells, and proliferative trophoblast cells were cultured as described above on glass-bottom multiwell plates (P24-1.5H-N, Cellvis). The cultures were fixed with 4% formaldehyde (28908, Thermo Fisher Scientific) in PBS for 20 min, followed by permeabilization with 0.2% Triton X-100 Surfact-Amps Detergent Solution (85111, Thermo Fisher Scientific) in PBS for 10 min. The only exception was sample preparation for *HLA-G* staining, which required fixation and permeabilization for 30 min with 100% methanol (322415, Millipore-Sigma) chilled to -20°C. Then the cultures were incubated with PBS supplemented with 0.2% bovine serum albumin (BSA; A9418, Millipore-Sigma) and 5% donkey serum (017-000-121, Jackson ImmunoResearch) for 1 h at RT, followed by incubation with primary antibodies overnight at 4°C. Secondary antibodies were incubated at 4°C for 2 h. Then the cultures were stained with 2 µM Hoechst 33342 (62249, Thermo Scientific) in PBS for 10 min before imaging on a Zeiss LSM 710 confocal microscope (most imaging experiments) or Leica DMi8 epifluorescent microscope (**xpression of ELF5 i, Extended Data Fig. 3**). Primary and secondary antibodies used are summarized in **Supplementary Table 8**.

### Bulk RNA-seq

hPSC, hPSC-derived TE cells, and proliferative TSC cultures were lysed using Buffer RLT Plus (1053393, Qiagen) supplemented with 2-mercaptoethanol (BME; 63689, Millipore-Sigma) directly in wells and RNA was extracted and purified using RNeasy Plus Mini Kit (74136, Qiagen) according to the manufacturer’s instruction. QIAcube Connect automated workstation was used for the extraction (Qiagen). Genomic DNA was eliminated by both the gDNA eliminator column and on-column incubation with DNase I (79256, Qiagen). RNA concentration and integrity was determined using RNA ScreenTape (5067-5576, Agilent Technologies) on the instrument 4200 TapeStation System (Agilent Technologies). All samples had the RNA Integrity Number (RIN) greater than 9.5. Fig. 1 sequencing libraries were constructed using TruSeq® Stranded mRNA Library Prep (20020595, Illumina) kit. Fig. 2 sequencing libraries were constructed using TruSeq® Stranded Total RNA Library Prep (20020597, Illumina) kit at the National Cancer Institute’s Center for Cancer Research sequencing core facility. The libraries were then sequenced at the same facility using the Illumina HiSeq system (Fig. 1) and the NovaSeq 6000 system (Fig. 2).

Bulk RNA-seq libraries for Figure 3 were constructed and sequenced in-house. In brief, 1 µg of purified RNA per sample was used to generate sequencing libraries following the manufacturer’s protocols of KAPA mRNA HyperPrep Kit (KK8581, Roche Life Science). The libraries were indexed using KAPA Unique Dual-Indexed Adapter Kit at 7 µM (KK8727, Roche Life Science) and 7 cycles of PCR library amplification were used. The sample preparation procedure was carried out using Biomek i7 Automated Workstation (Beckman Coulter) and the automation protocol was validated by Roche Life Science. Libraries were then quantified on a QuantStudio Real-Time PCR System (A34322, Applied Biosystems) using the KAPA Library Quantification Kit for Illumina Platforms (KK4824, Roche Life Science). The libraries were individually normalized to 4 nM by diluting each library with 10 mM Tris-HCl (pH 8.5; T1062, Teknova) prior to pooling. The pooled libraries were then quantified according to the manufacturer’s instruction and diluted to 1.5 nM for sequencing. Sequencing was performed on Illumina NovaSeq 6000 system using NovaSeq 6000 S4 Reagent Kit v1, 300 cycle (20012866, Illumina), with 300 pM as the final loading concentration.

### Bulk RNA-seq data analysis

The details of the data analysis procedures are described in Supplementary Methods. Analysis scripts are available at: https://github.com/cemalley/Slamecka_methods.

### PluriTest

FASTQ files corresponding to RNA-seq samples were first downsampled so that the file sizes did not exceed 1.5 GB, a prerequisite for the web-based analysis. The FASTQ files were then uploaded to https://www.pluritest.org for analysis in pair-end mode.

### scRNA-seq

Single-cell suspensions from hPSC, D3, D6 TE cells, and TSC were obtained after a 15 min Accutase treatment; and from D9 TE cells after 20-25 min. Cells were suspended in PBS+0.04% BSA and cell clumps were mitigated by passing the suspension through the cell strainer pluriStrainer® with pore size of 20 µm (43-50020-03, pluriSelect). Gel Bead-In Emulsion (GEM) generation, cDNA synthesis, and sequencing library preparation was performed in-house using Chromium Single Cell 3’ Library & Gel Bead Kit version 2 (120237, 10X Genomics) and Chromium Single Cell A Chip Kit (120236, 10X Genomics). Figure 3 single-cell sequencing libraries were prepared using Chromium Single Cell 3’ GEM, Library & Gel Bead Kit version 3 (1000075, 10X Genomics) and Chromium Single Cell B Chip Kit (1000073, 10X Genomics). Sample indexing for both scRNA-seq experiments was performed using Chromium i7 Multiplex Kit (120262, 10X Genomics). To determine the concentration, integrity, and size distribution of fragments during the procedure, cDNA trace analysis was performed using High-Sensitivity D5000 ScreenTape (5067-5588, Agilent Technologies) and library trace analysis was performed using High-Sensitivity D100 ScreenTape (5067-5582, Agilent Technologies) on the 4200 TapeStation System (Agilent Technologies). The prepared libraries for the Fig. 2 dataset were sequenced using the NovaSeq 6000 at the National Cancer Institute’s Center for Cancer Research sequencing core facility. The Fig. 3 dataset single-cell sequencing libraries were sequenced in-house using the NovaSeq 6000. scRNA-seq parameters are summarized in **Supplementary Table 9** and include sample indices, numbers of cDNA reverse transcription (RT) cycles, and numbers of targeted and recovered cells.

### scRNA-seq data analysis

The details of the data analysis procedures are described in Supplementary Methods. Analysis scripts are available at: https://github.com/cemalley/Slamecka_methods.

### miRNA-seq

Total RNA of each sample was extracted from cell pellets using QIAzol Lysis Reagent (79306, Qiagen) and used to prepare the miRNA sequencing library. The assay and a part of the data analysis was performed by Arraystar Inc. After the completed libraries were quantified with the Agilent 2100 Bioanalyzer, the DNA fragments in the libraries were denatured with 0.1M NaOH (72068, Millipore-Sigma) to generate single-stranded DNA molecules. The single-stranded DNA molecules were then captured on Illumina flow cells, amplified *in situ,* and sequenced for 51 cycles on Illumina NextSeq 500 according to the manufacturer’s instruction. Details of the method and analysis are described in Supplementary Methods.

### MeDIP-seq

The assay and a part of the data analysis was performed by Active Motif. The samples were delivered to Active Motif as flash-frozen cell pellets on dry ice. DNA was extracted, then sonicated to ∼150-300 bp, and Illumina adapters were ligated to the DNA ends. This DNA was then used in immunoprecipitation (IP) reactions using 5-Methylcytosine (5-mC) mouse monoclonal antibody (39649, Active Motif). Immunoprecipitated DNA and input control (pooled DNA that did not go through the IP step) were finally processed into sequencing libraries using PrepX DNA Library Kit (400075, Takara) and sequenced using the Illumina platform (NextSeq 500, 75-nt single-end). Details of the data analysis are described in Supplementary Methods.

### Periodic Acid-Schiff (PAS) staining

PAS staining was performed using the Periodic Acid-Schiff (PAS) Kit (395B, Millipore-Sigma). TE and TSC cultures were grown on VTN-coated 6-well plates in their respective media, with daily medium change, as described above. Then the cultures were fixed for 1 min with a solution of 28.8% formaldehyde and 10% ethanol (E7023, Millipore-Sigma) in water. The fixed cultures were washed with slowly running tap water for 1 min, treated with periodic acid for 5 min, and washed three times with distilled water. The cells were then treated with Schiff reagent for 15 min, washed with running tap water for 5 min, and then imaged using a Zeiss Axiovert microscope equipped with an Axiocam 506 color camera with no phase contrast applied.

### TEM

hPSC differentiating into TE at different time points were fixed with glutaraldehyde in cacodylate buffer prepared by the Electron Microscopy Core at the Center for Cancer Research (National Cancer Institute at Frederick) for 24 h and further processed and imaged at the same facility.

### Pregnancy Test

The spent medium of TE cells differentiated from hPSC at D10 (cultured in TE2 medium) was loaded into the cassette of a commercially available over-the-counter Alere hCG Urine II Test Kit for detection of hCG. The hCG detection threshold is 20 mIU/mL according to the manufacturer (Abbott).

### STB fusion time-lapse imaging

hPSC were seeded into VTN-coated Incucyte ImageLock 96-well plates and differentiated into TE. On day 5, the plates were transferred into an Incucyte S3. Time-lapse phase-contrast were taken every 4 h for 14 additional days.

### Western blot analysis

The automated, capillary-based Western blotting system Wes (ProteinSimple) was used according to the manufacturer’s instructions. Briefly, cells were lysed in RIPA Lysis and Extraction Buffer (89901, Thermo Fisher Scientific) supplemented with cOmplete™, Mini Protease Inhibitor Cocktail (04-693-124-001, Roche), assisted by sonication. Lysates were cleared of debris by centrifugation at 14,000 g for 15 min. The protein content in lysates was determined using Pierce™ BCA Protein Assay Kit (23225, Thermo Fisher Scientific) according to the manufacturer’s instructions. Lysates were diluted 1:4 with 1× sample buffer (ProteinSimple). The capillary cartridges of the 12-230 kDa Separation Module (SM-W003, ProteinSimple) were used, along with Anti-Rabbit (DM-001, ProteinSimple) and Anti-Mouse (DM-002, ProteinSimple) Detection Modules containing reagents and HRP-conjugated secondary antibodies. The detected chemiluminescent signal data were analyzed using the ProteinSimple Compass software. All western blot data were displayed by lanes in virtual blot-like images. Primary antibodies used are listed in **Supplementary Table 8.**

### qRT-PCR

Cultures of human dermal fibroblasts, JEG-3 CC cells, and hPSC-derived TSCs were grown in 6-well plates. Cell culture medium was aspirated from the wells, the cells were washed twice with PBS, and the cell monolayers were lysed with Buffer RLT Plus supplemented with BME. RNA was extracted and purified using an RNeasy Plus Mini Kit (74136, Qiagen) according to the manufacturer’s instructions. cDNA was synthesized from 500 ng of total RNA using a High-Capacity RNA-to-cDNA™ Kit (4388950, Thermo Fisher Scientific). PrimeTime Std® qPCR Assays (Integrated DNA Technologies) were used, according to manufacturer’s instructions, as sets of primers and probes against gene targets. The instrument used was QuantStudio 12K Flex Real-Time PCR System (Thermo Fisher Scientific). Each reaction was performed in a final volume of 10 µl, consisting of 5 µl of 2× PrimeTime® Gene Expression Master Mix (1055771, Integrated DNA Technologies), 0.5 µl of 20× primer/probe suspension and 4.5 µl of cDNA diluted in DNase/RNase-free water (10977015, Thermo Fisher Scientific). The thermocycler program consisted of an initial UDG incubation at 50°C for 2 min, enzyme activation at 95°C for 10min, followed by 40 cycles at 95°C for 15 s and 60°C for 30 s. To confirm product specificity, melting curve analysis was performed after each amplification. GAPDH was used as a housekeeping gene. List of primer-probes is provided in **Supplementary Table 9.**

### Secretome analysis

WA09 ESCs (D4) and WA09-derived TE cells (D10) were grown in VTN-coated 6-well plates at 37°C in a humidified atmosphere containing 5% CO2 and 21% O2. A total of 8 ml of spent medium was collected from 4 wells of a 6-well plate after 24 h. The medium was then frozen at -80°C and shipped to Applied Biomics for analysis using the nanoscale liquid chromatography coupled to tandem mass spectrometry (Nano LC-MS/MS) method. The main steps performed were protein fractionation, reduction, alkylation, trypsin digestion, and Nano HPLC. Ion composition was then detected by MS/MS and a database search was then performed to identify the proteins.

### Calcein AM and propidium iodide staining

TSCs were seeded onto VTN-coated tissue culture-treated 12-well plates at a density of 4 × 10^5^/cm^2^ and cultured in TE3 medium. 24 h following seeding, 2 µM Calcein AM, a component of the LIVE/DEAD™ Viability/Cytotoxicity Kit for mammalian cells (L3224, Thermo Fisher Scientific), and 1µg/ml propidium iodide (J66584-AB, Alfa Aesar) were added into each well without medium change to fluorescently mark viable and dead cells, respectively. Incucyte S3 was used for live-cell imaging of the cultures 30 min following staining and for image analysis. The percentage of dead cells out of all cells was then calculated.

### Data and analysis code availability

All sequencing data was deposited to the NCBI Short Read Archive (SRA) under accession PRJNA760795.

Analysis code is available at https://github.com/cemalley/Slamecka_methods.git.

