## Supplementary figures and images for "A Comprehensive Roadmap of Human Placental Development *in vitro*"

### Ext Data Fig 1

**Extended Data Figure 1 (Slamecka et al.)**

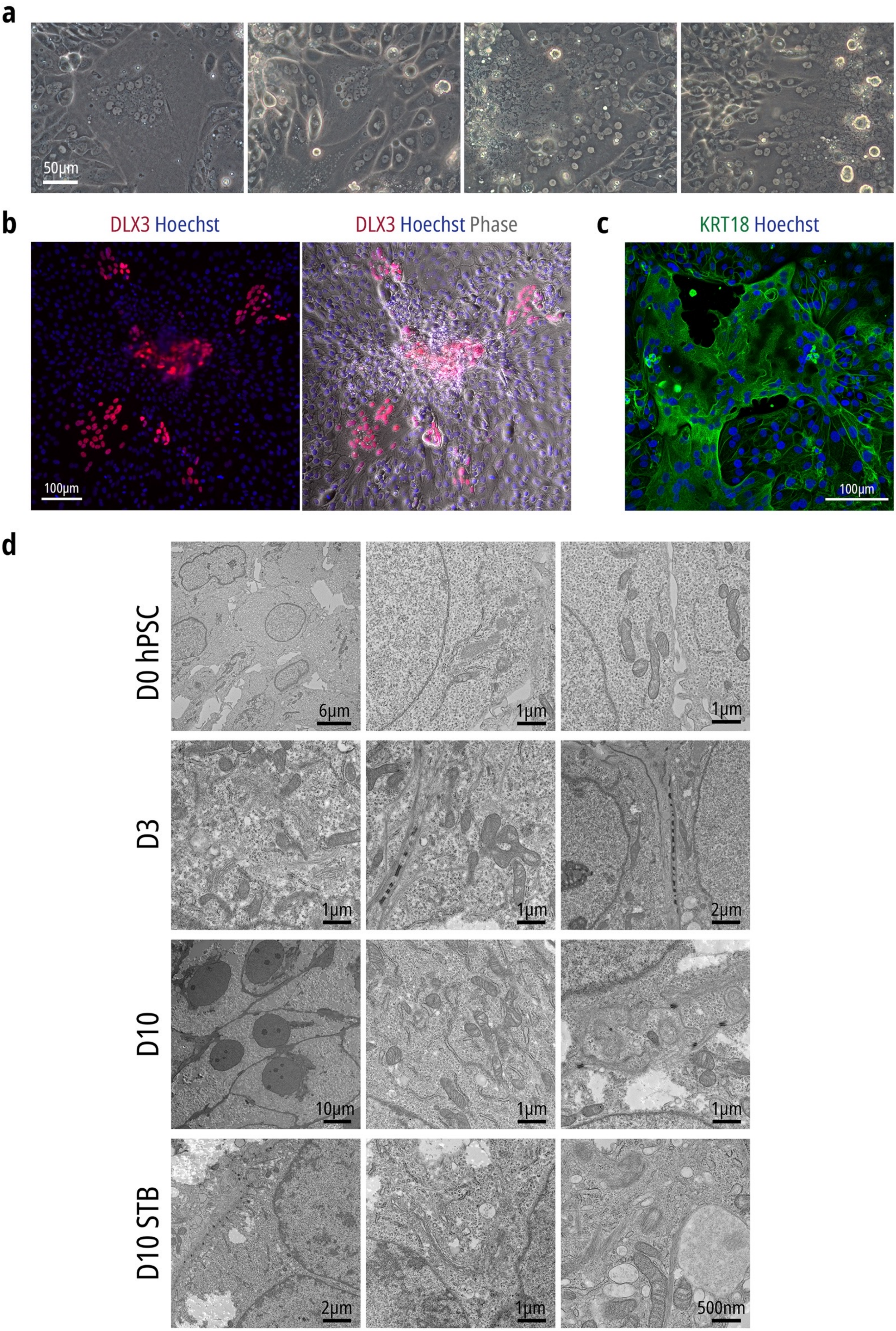

### Ext Data Fig 2

#### Extended Data Figure 2 (Slamecka et al.)

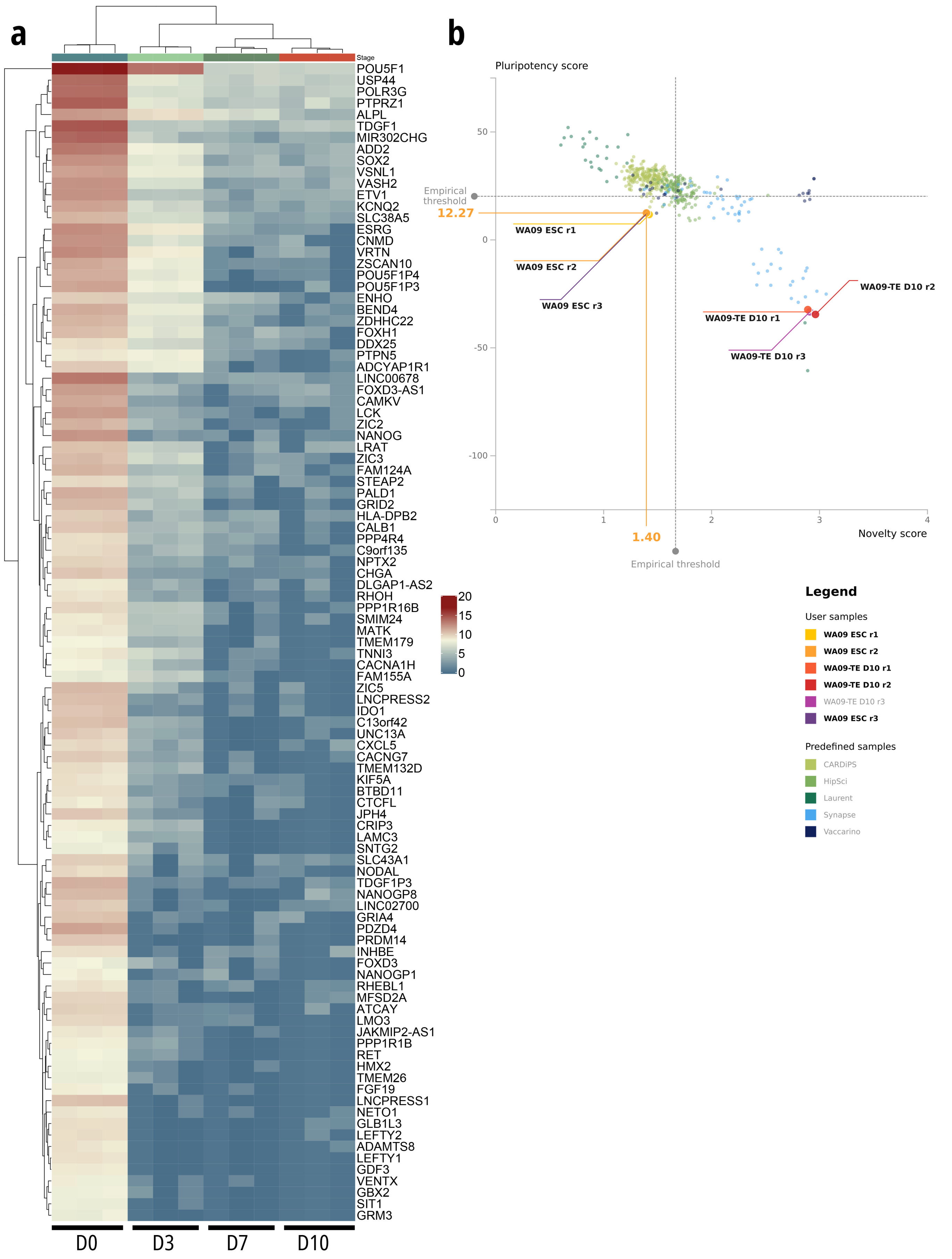

### Ext Data Fig 3

**Extended Data Figure 3 (Slamecka et al.)**

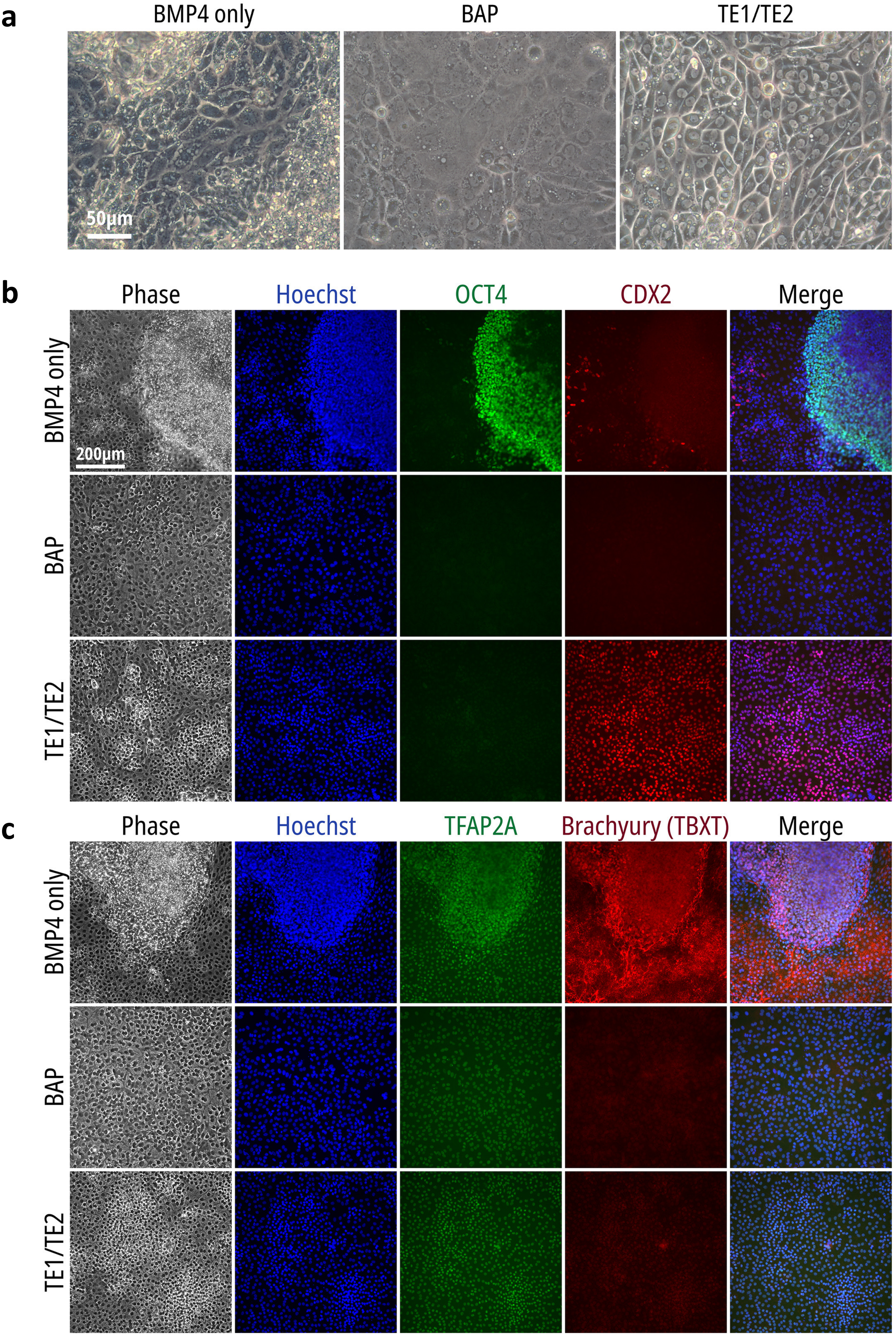

### Ext Data Fig 4

#### Extended Data Figure 4 (Slamecka et al.)

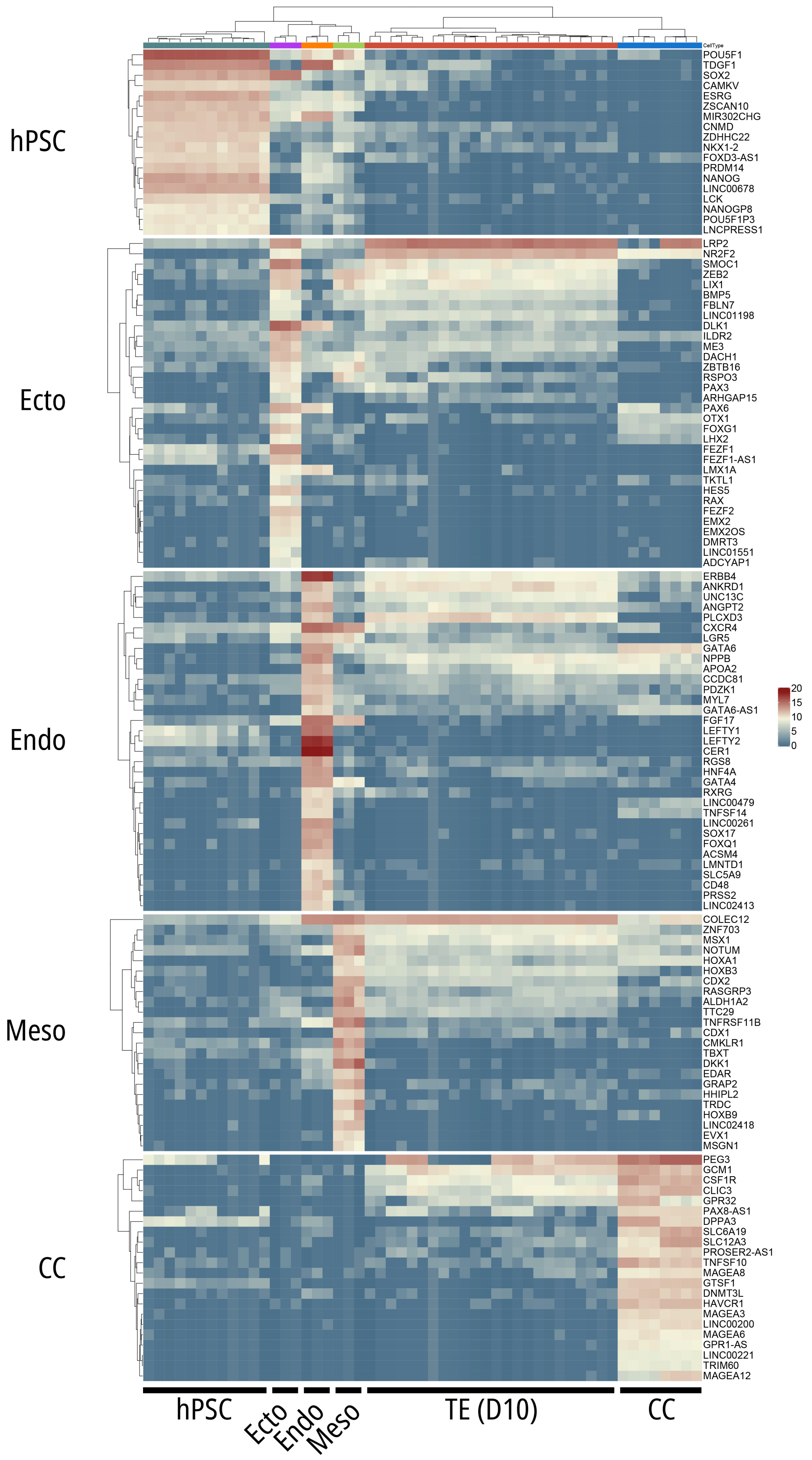

### Ext Data Fig 5

**Extended Data Figure 5 (Slamecka et al.)**

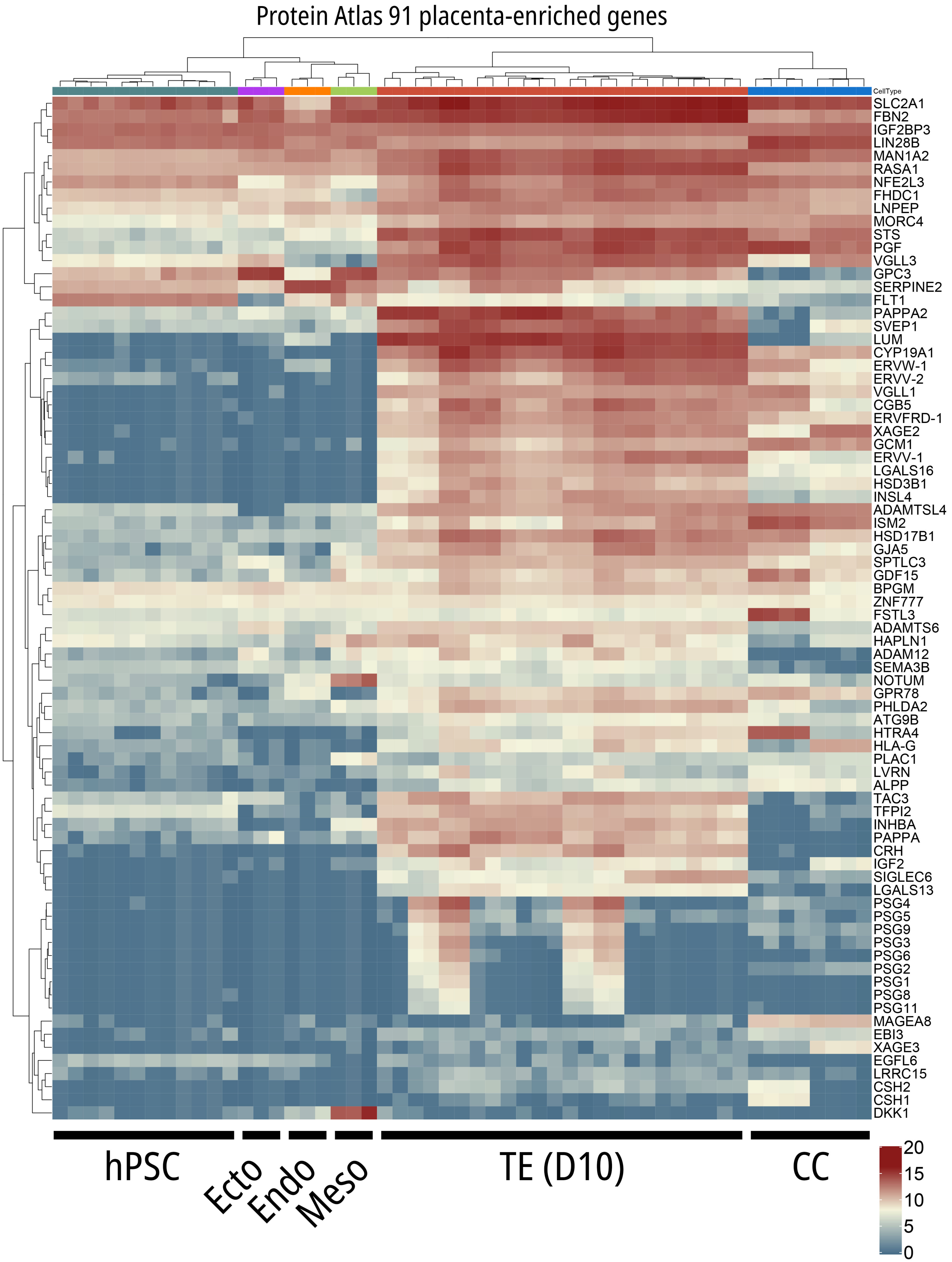

### Ext Data Fig 6

**Extended Data Figure 6 (Slamecka et al.)**

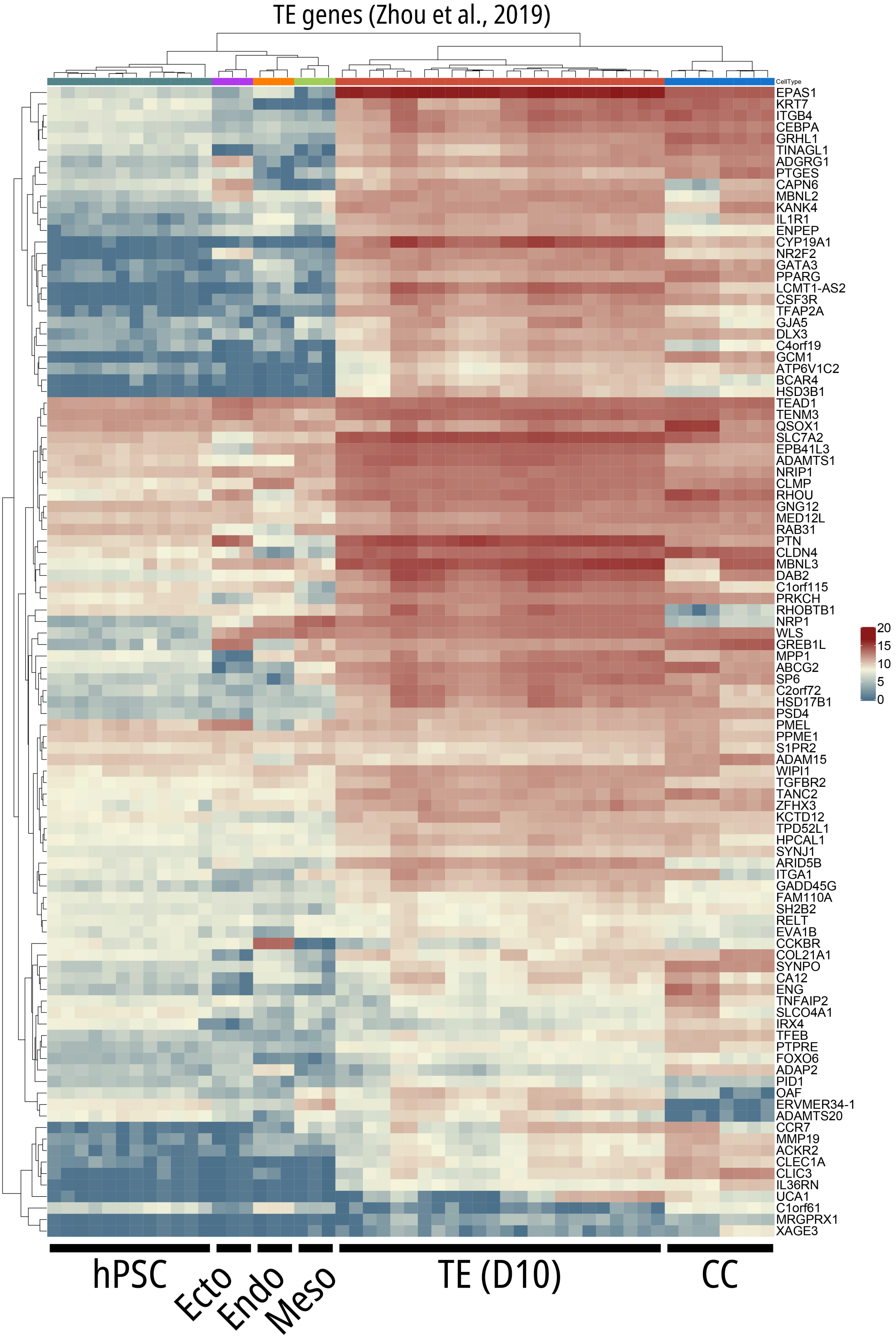

### Ext Data Fig 7

#### Extended Data Figure 7 (Slamecka et al.)

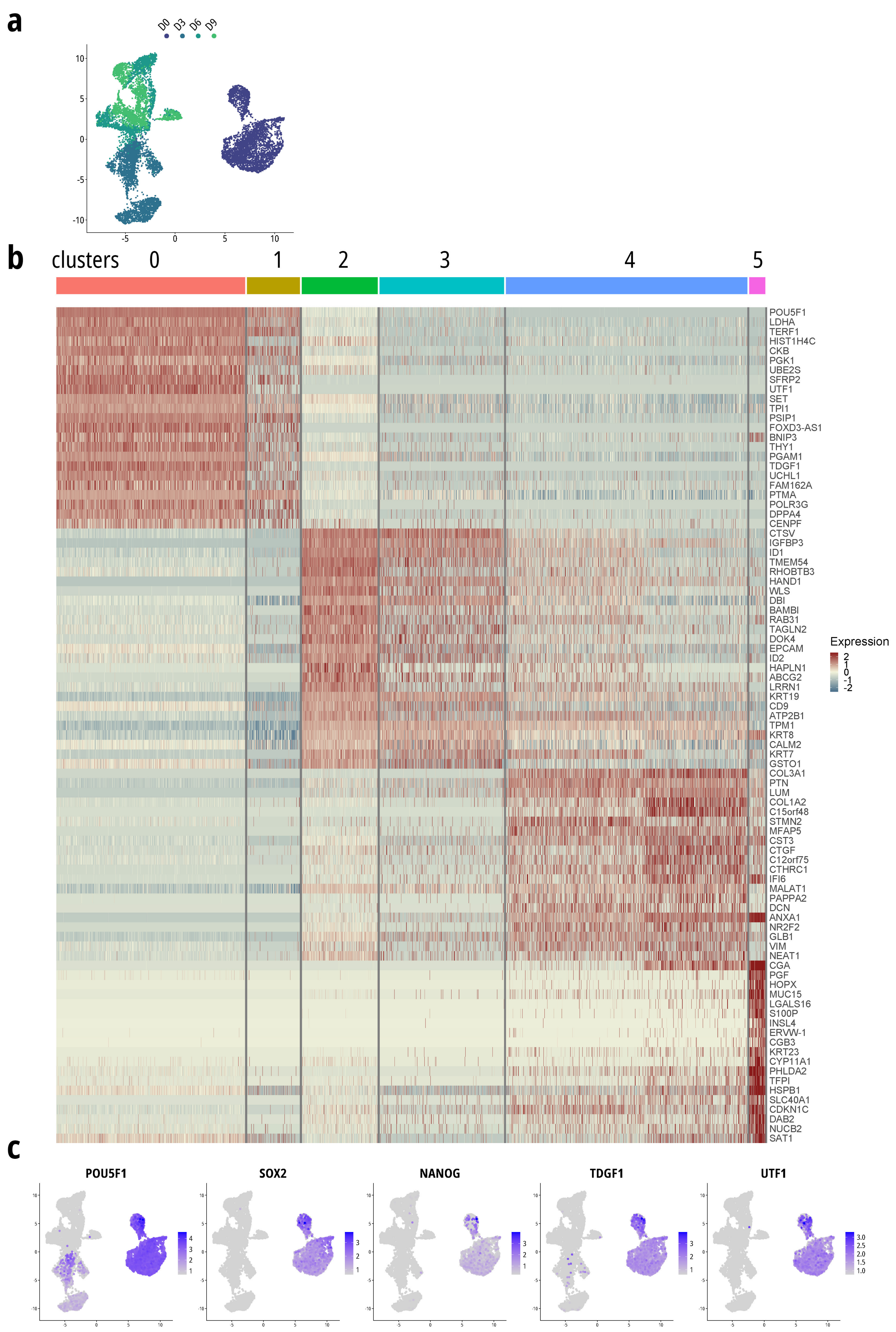

### Ext Data Fig 8

**Extended Data Figure 8 (Slamecka et al.)**

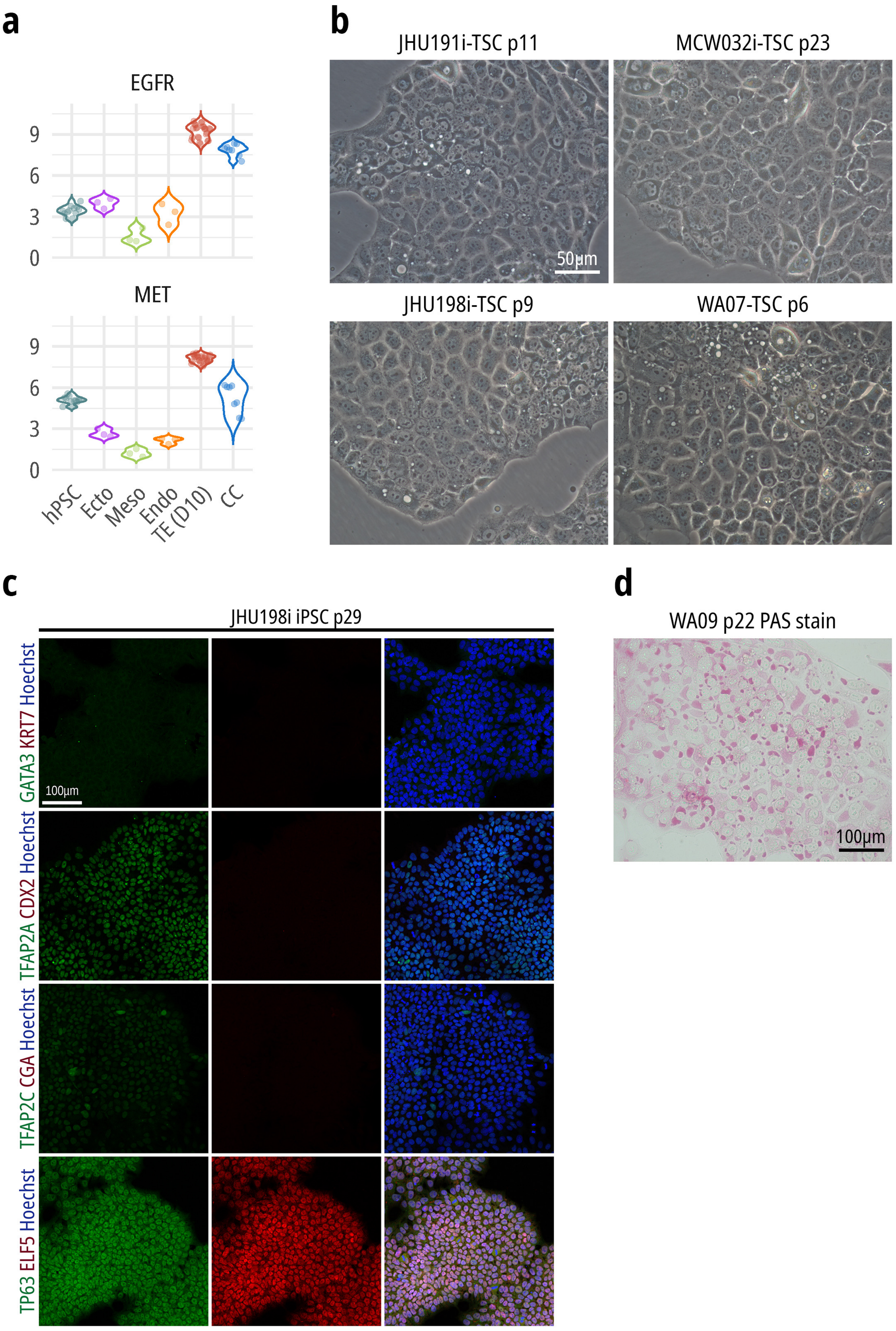

### Ext Data Fig 9

**Extended Data Figure 9 (Slamecka et al.)**

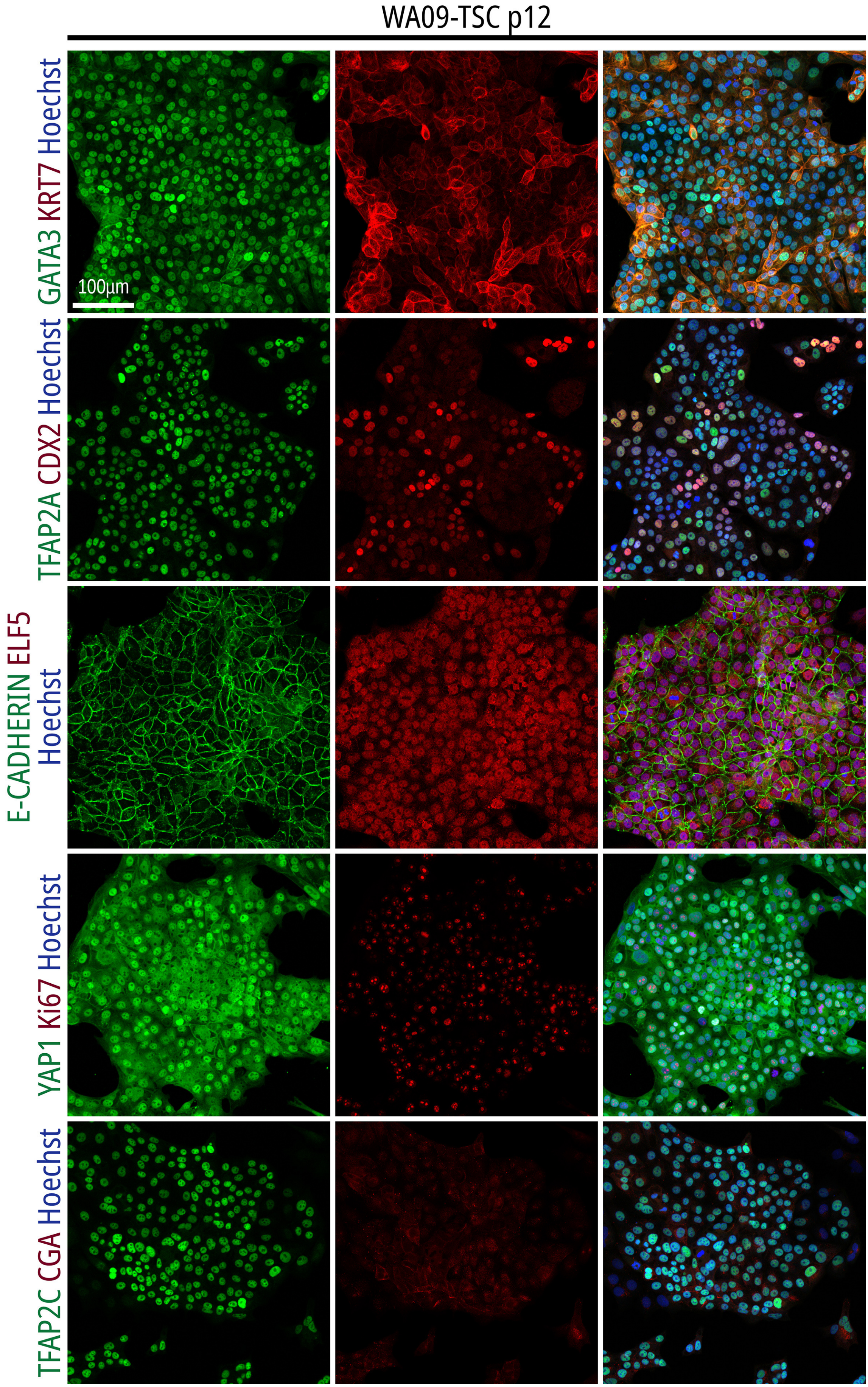

### Ext Data Fig 10

Extended Data Figure 10 (Slamecka et al.)

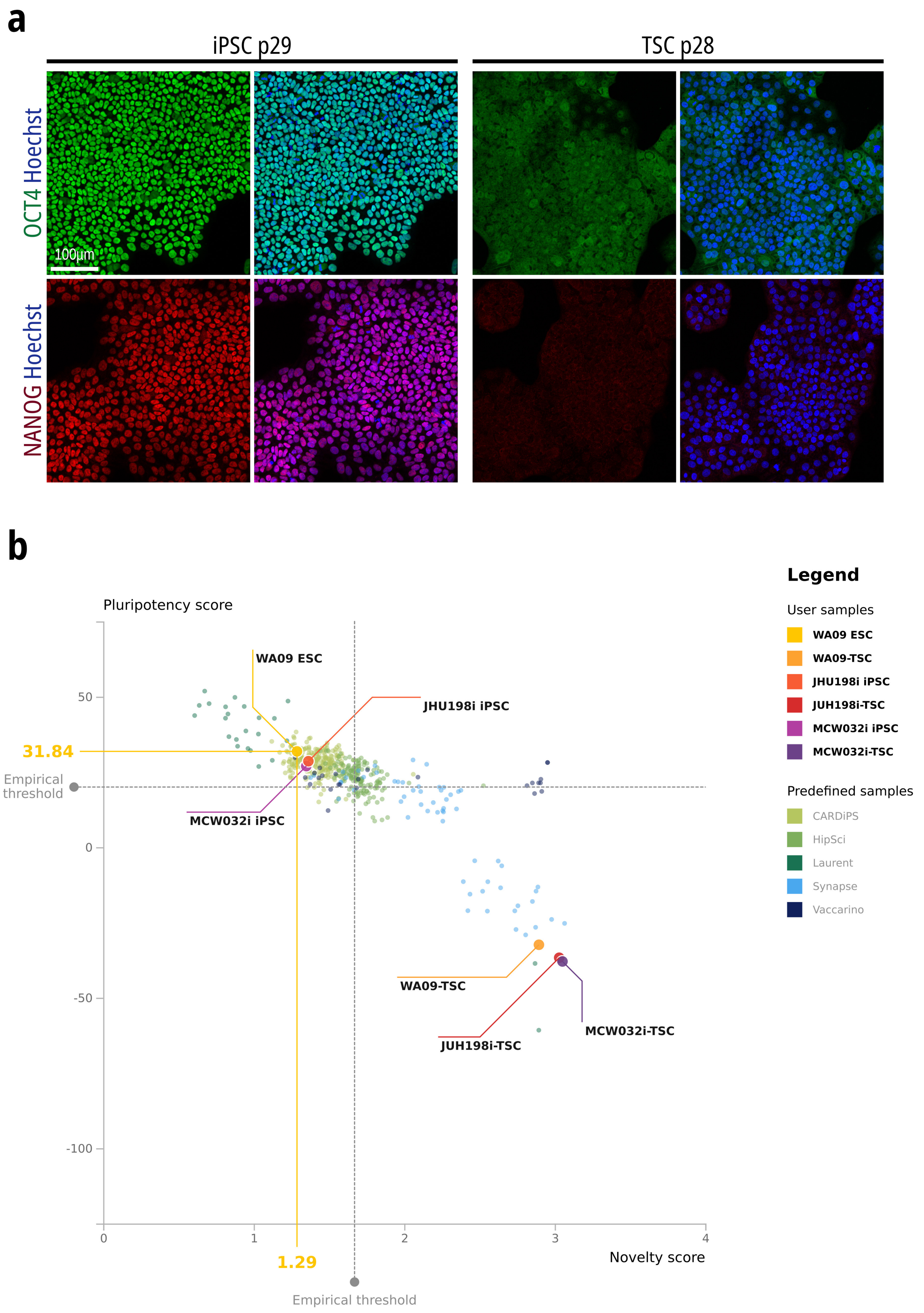

### Ext Data Fig 12

**Extended Data Figure 12 (Slamecka et al.)**

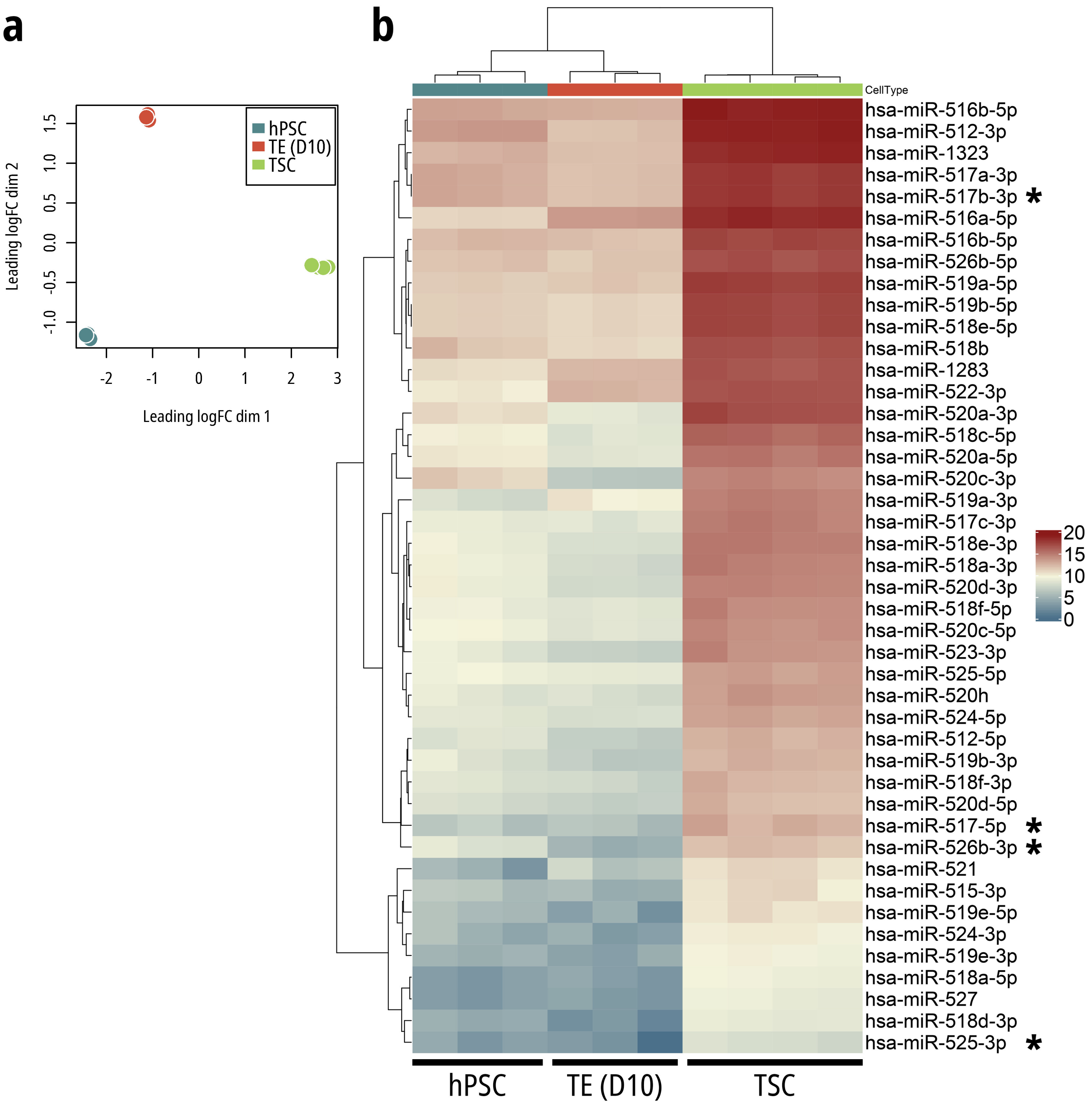

### Ext Data Fig 13

#### Extended Data Figure 13 (Slamecka et al.)

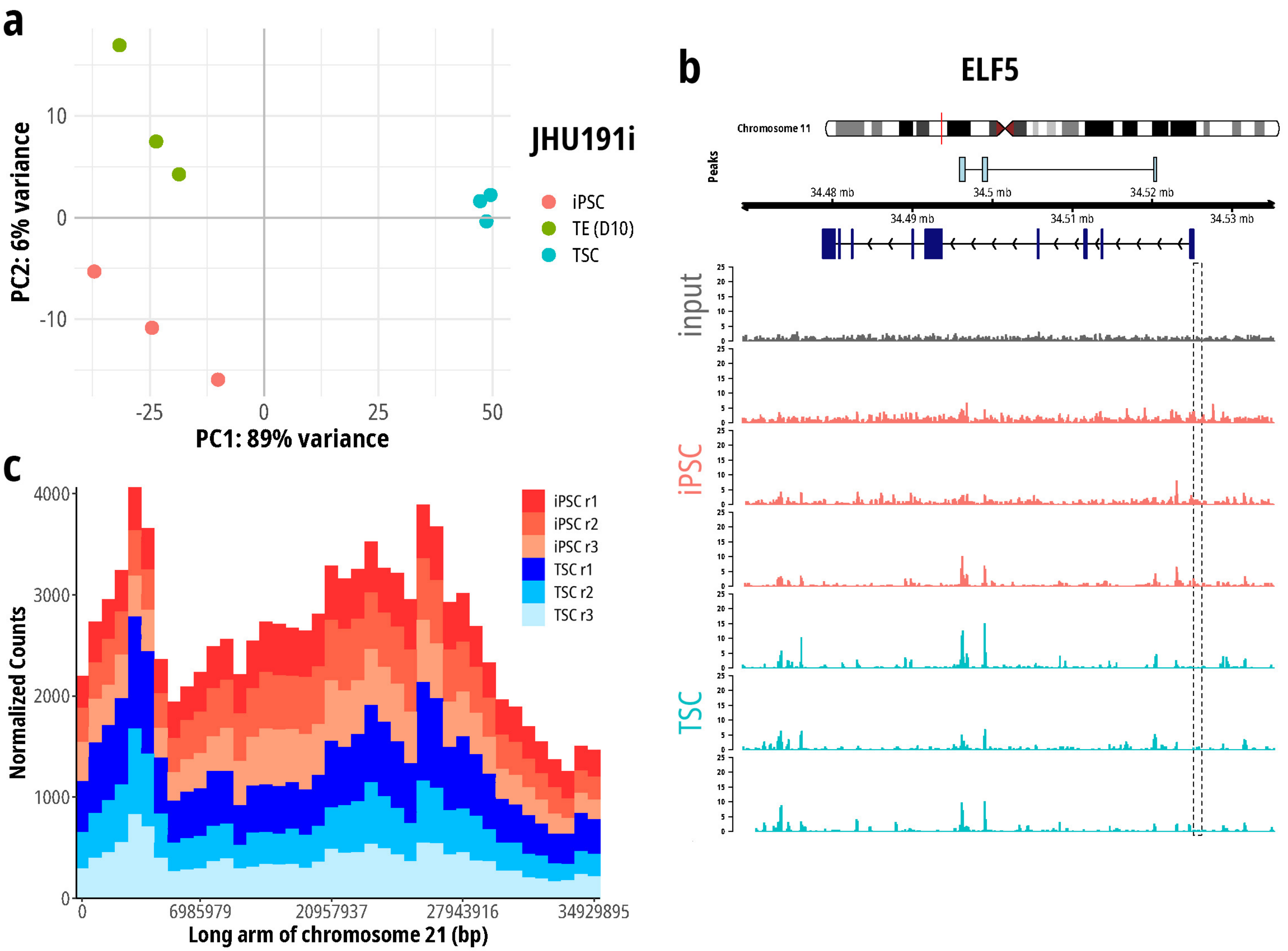

### Ext Data Fig 14

#### Extended Data Figure 14 (Slamecka et al.)

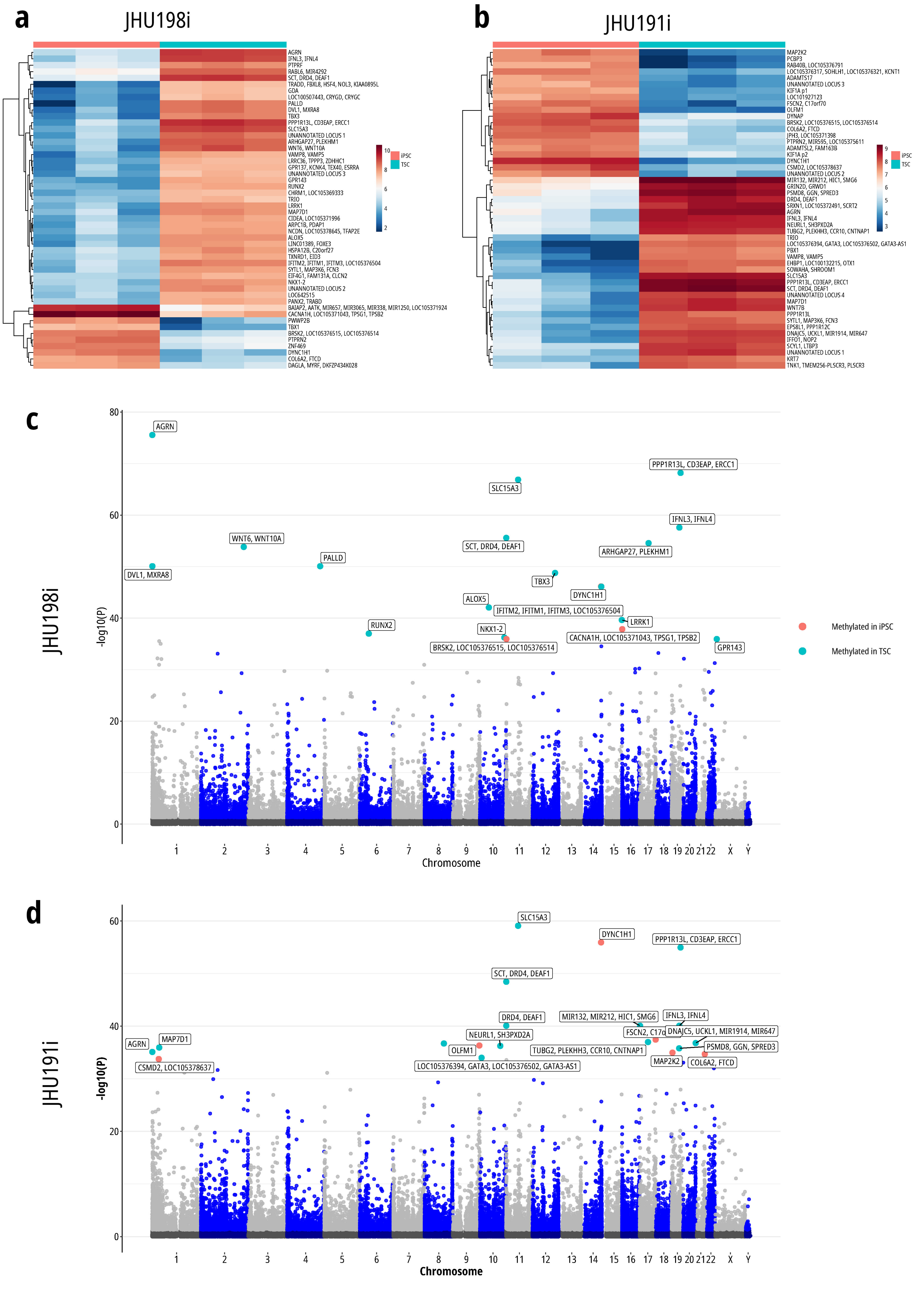

### Ext Data Fig 15

#### Extended Data Figure 15 (Slamecka et al.)

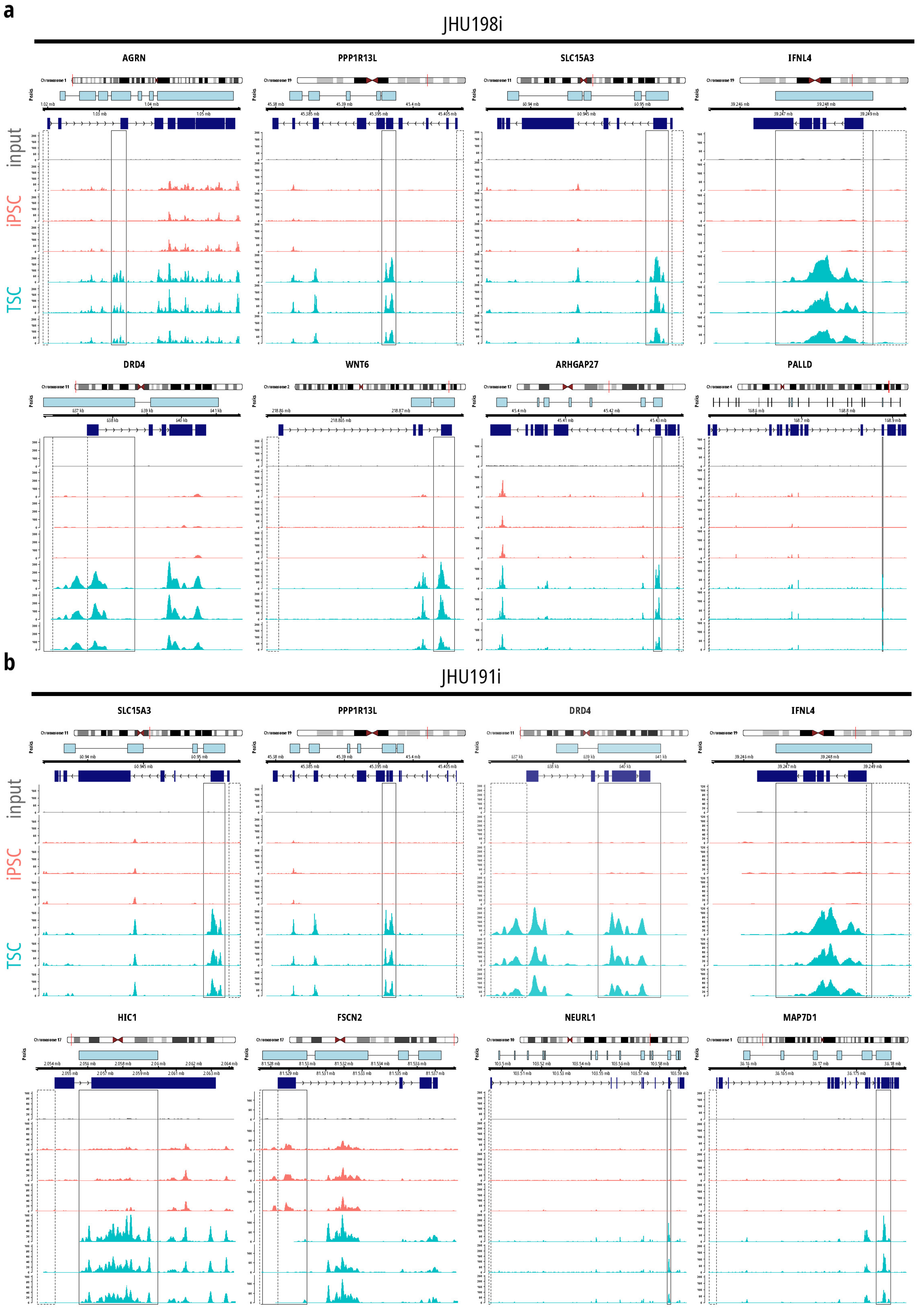

### Ext Data Fig 16

**Extended Data Figure 16 (Slamecka et al.)**

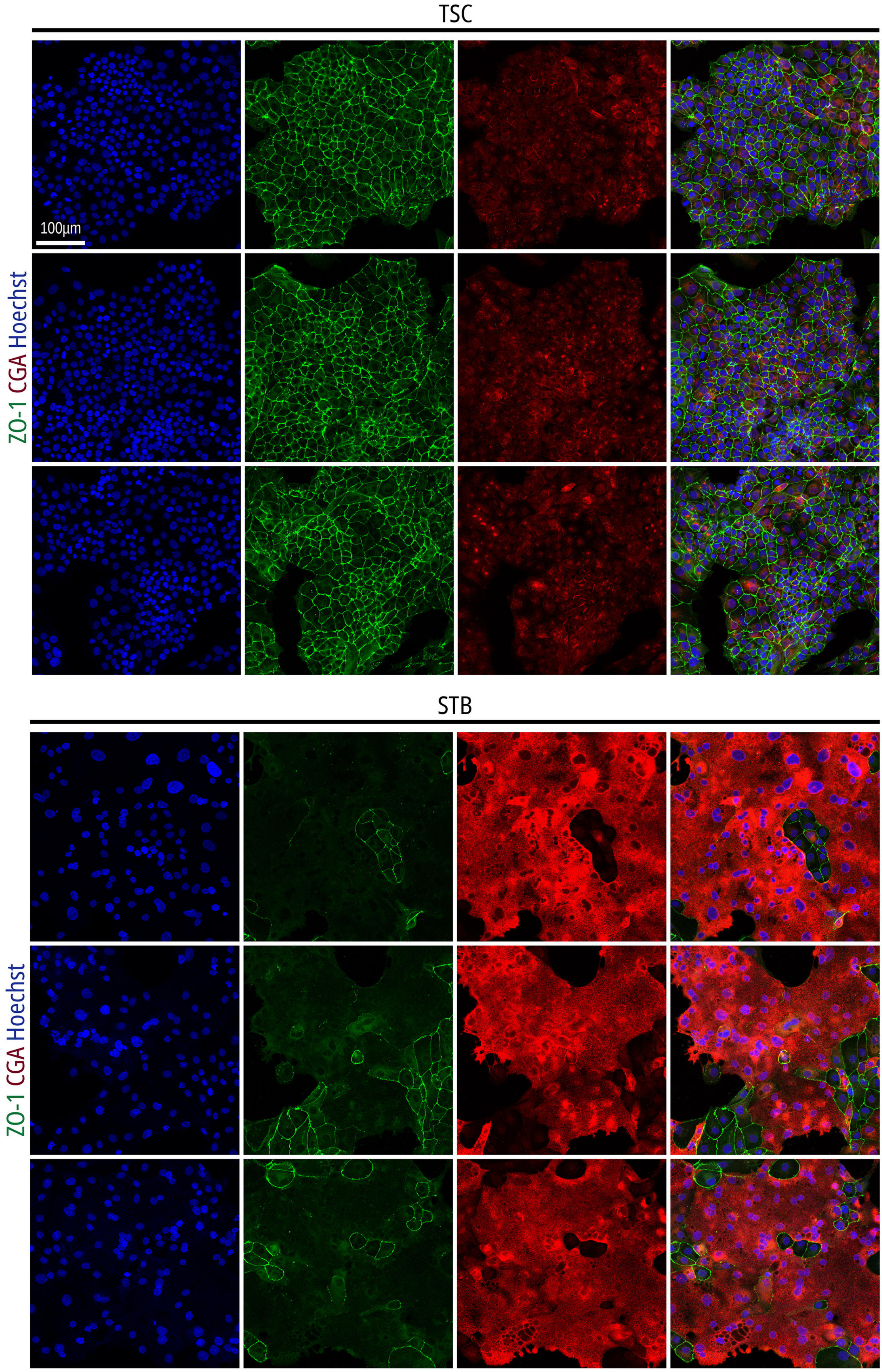

### Ext Data Fig 17

**Extended Data Figure 17 (Slamecka et al.)**

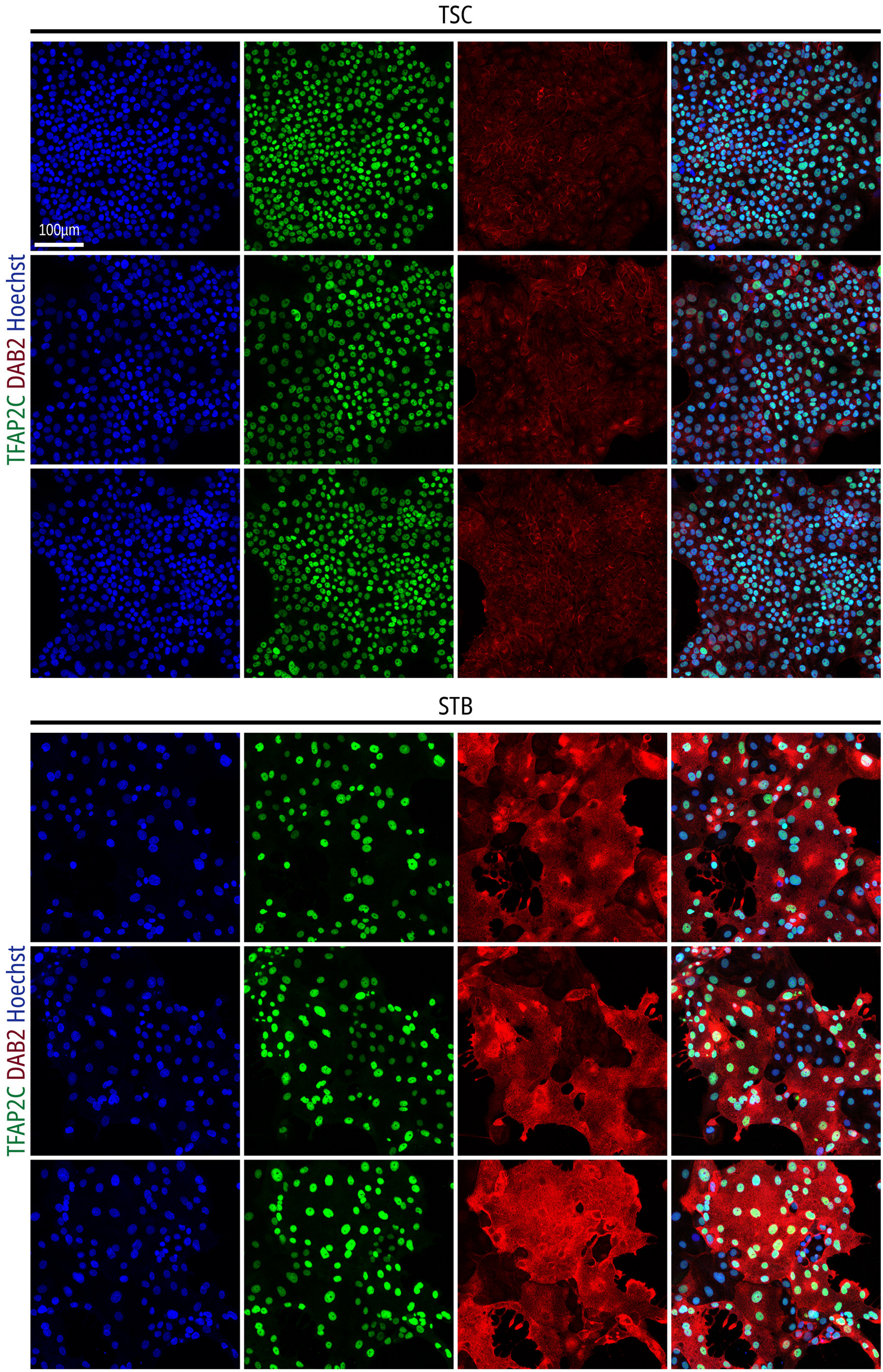

### Ext Data Fig 18

##### Extended Data Figure 18 (Slamecka et al.)

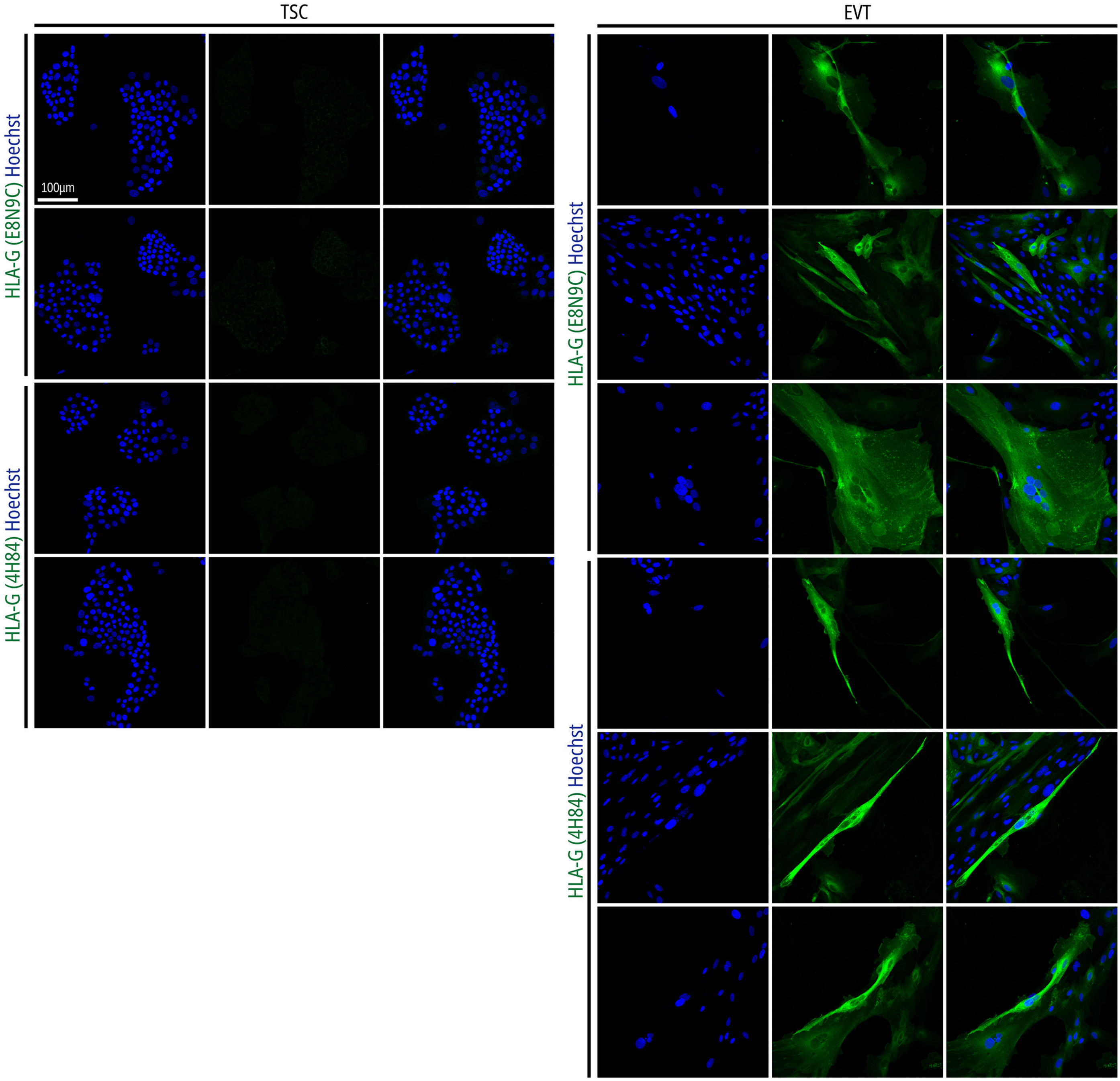

### Ext Data Fig 19

**Extended Data Figure 19 (Slamecka et al.)**

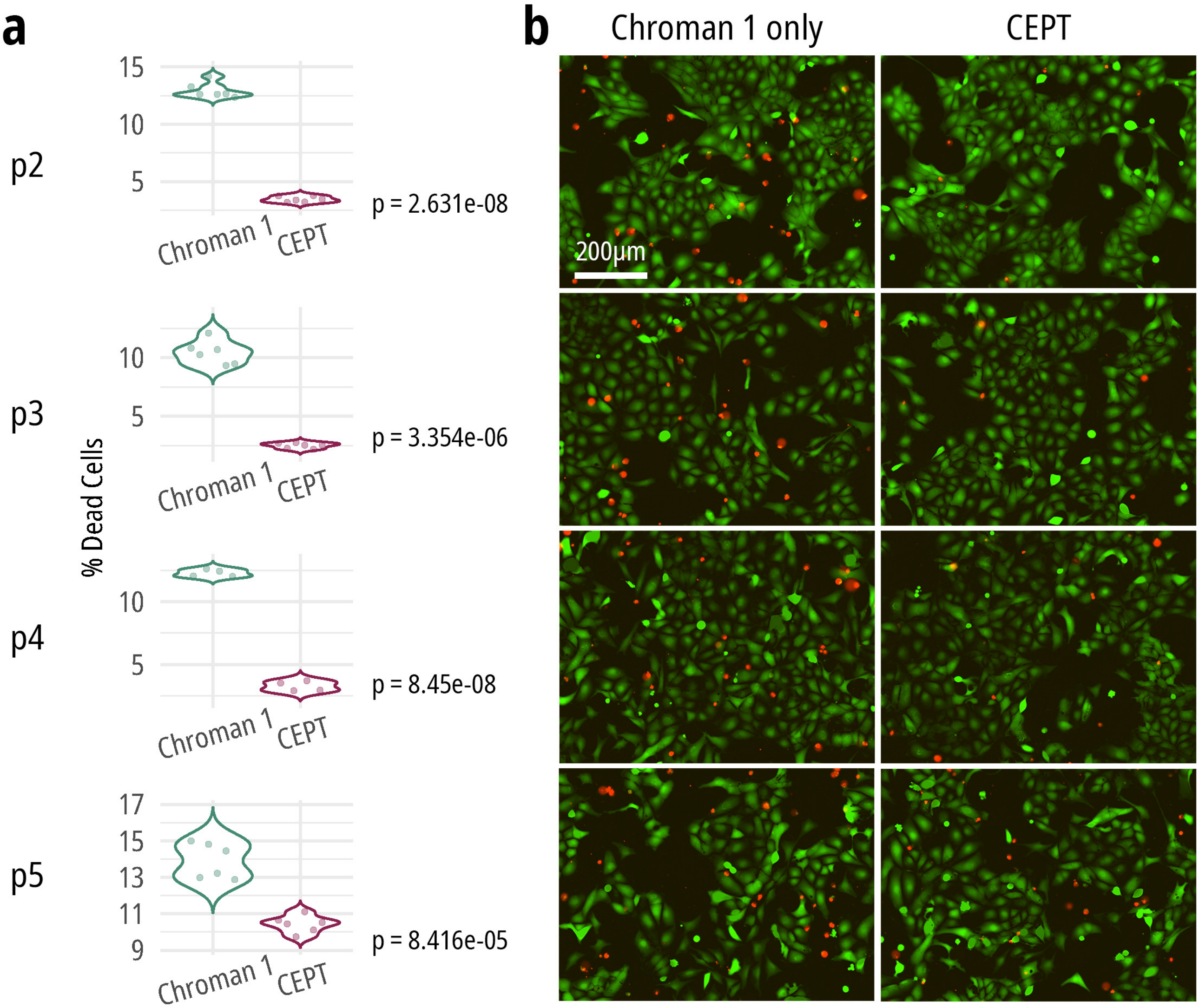

### Ext Data Fig 20

**Extended Data Figure 20 (Slamecka et al.)**

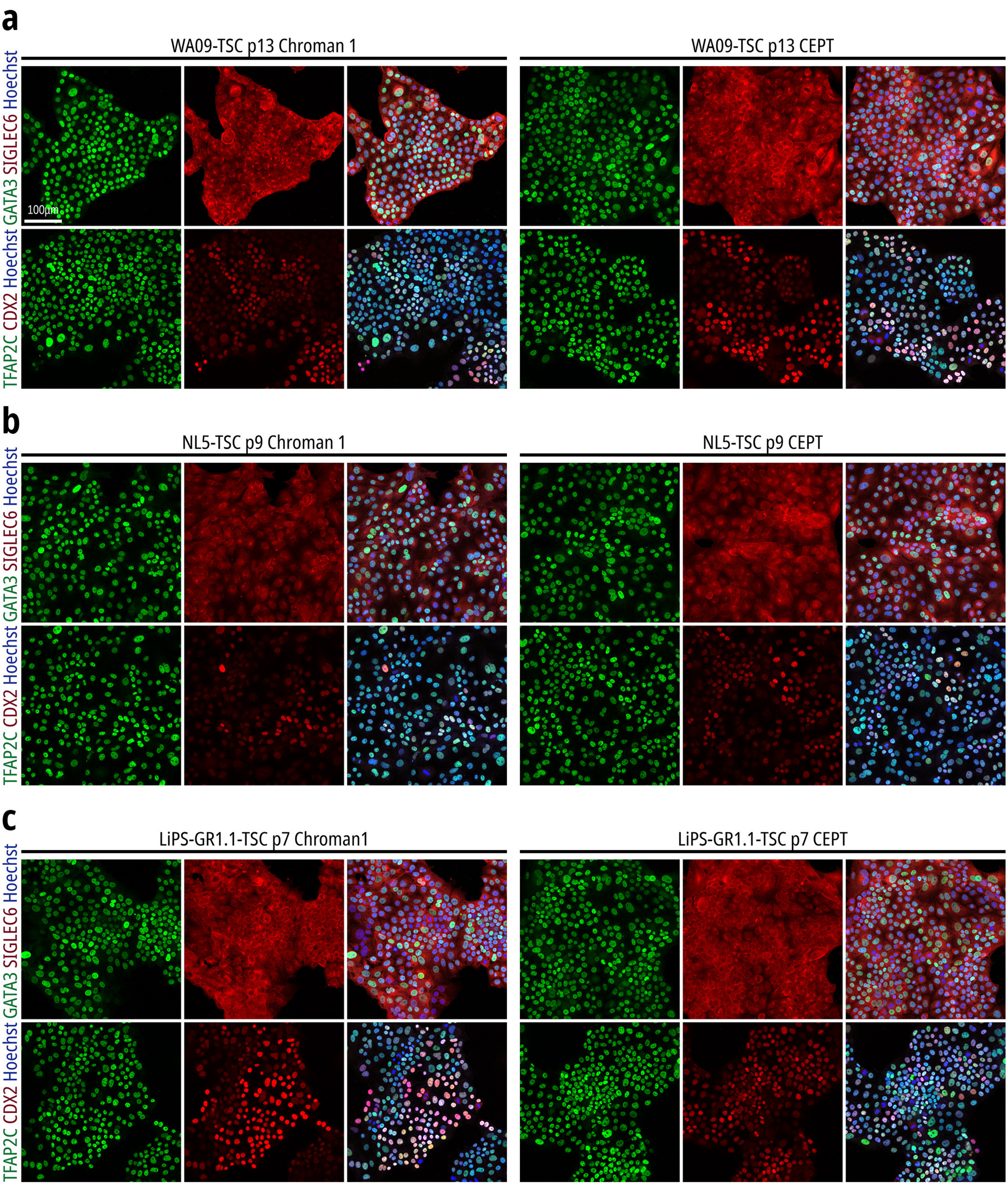
