## Supplementary material for "A Comprehensive Roadmap of Human Placental Development *in vitro*": Ext Data Fig 11

### Extended Data Figure 11 (Slamecka et al.)

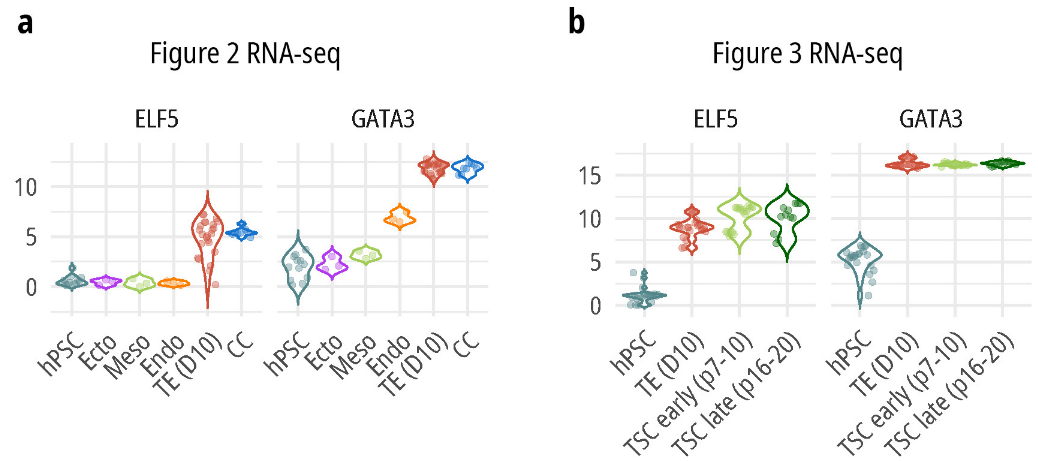


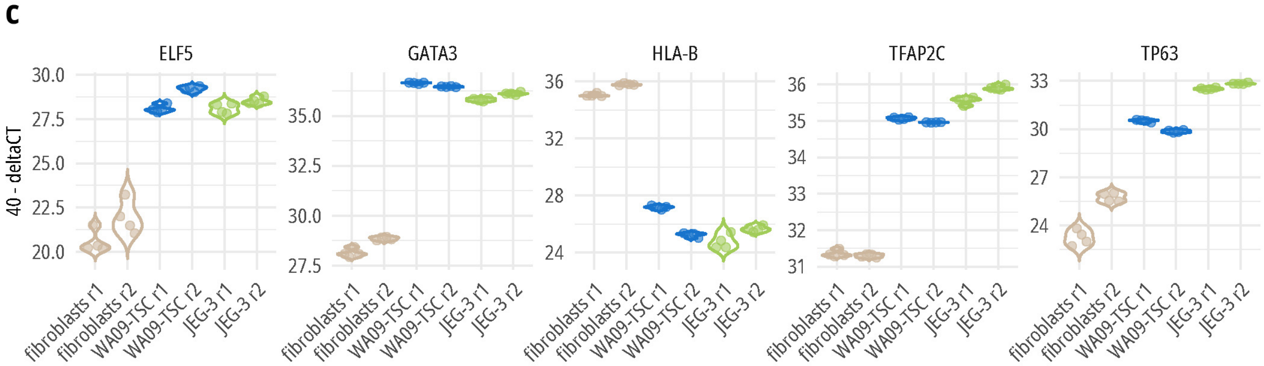


| | logFC TSC vs. fibroblasts | adj. P value | logFC JEG-3 vs. fibroblasts | adj. P value | logFC TSC vs. JEG-3 | adj. P value |
|---|---|---|---|---|---|---|
| **GATA3** | 8.07 | 1.3E-25 | 7.48 | 3.54E-25 | 0.59 | 3.81E-04 |
| **TFAP2C** | 3.69 | 3.17E-24 | 4.41 | 1.19E-25 | -0.72 | 6.89E-09 |
| **HLA-B** | -9.22 | 5.37E-18 | -10.22 | 5.65E-19 | 1.00 | 0.01 |
| **ELF5** | 7.40 | 4.11E-16 | 7.06 | 9E-16 | 0.34 | 0.35 |
| **TP63** | 5.69 | 1.35E-12 | 8.18 | 8.62E-16 | -2.49 | 8E-06 |
