## Supplementary Methods for "A Comprehensive Roadmap of Human Placental Development *in vitro*"

Jaroslav Slamecka, Carlos A. Tristan, Seung Mi Ryu, Pei-Hsuan Chu, Claire Weber, Tao Deng, Yeliz Gedik, Pinar Ormanoglu, Sam Michael, Ty C. Voss, Anton Simeonov, Ilyas Singeç

### Bulk RNA sequencing data analysis

Raw FASTQ files were aligned to the human reference genome^1^ (CRCh38 primary assembly, Ensembl annotation version 100) using STAR^2^ (version 2.7.5a). Counts were derived from the aligned reads using featureCounts from the Subread suite^3^ (v 2.0.0). The counts were processed in R^4^ (version 4.0.3) using package edgeR (version 3.34.0)^5^. Filtering of genes with low counts, normalization, multidimensional scaling (MDS) plot construction and Differential expression analysis were performed using limma (version 3.48.1)^6^. Heatmaps were constructed using ComplexHeatmap (version 2.8.0)^7^. Expression plots were constructed using ggplot2 (v3.3.5)^8^. Gene set enrichment analysis was performed in R using Enrichr (package “enrichR”)^9–11^. The analysis scripts are available at <https://github.com/cemalley/Slamecka_methods>.

All computationally intensive operations for all analyses presented in this manuscript were performed on the NIH’s Linux-based high-performance computing platform Biowulf.

### Bulk RNA-seq integration analysis

For the purpose of comparison of RNA-seq transcriptional profiles obtained as part of this study and external laboratories, raw data in form of FASTQ files was downloaded from the Sequence Read Archive (SRA)^12^ using the command fasterq-dump from the SRA toolkit (v2.11.0). The mapping/alignment and feature counting were performed using the same workflow, the same versions of reference genome and software versions, as were used for the datasets presented in this study. Then the counts were collectively loaded into R using edgeR and the batch effect was removed using ComBat-seq, applying it to the raw, unfiltered counts by supplying the batch variable. No group variable was specified. Then the counts were filtered, normalized, and MDS plots were constructed using limma.

### Single-cell RNA sequencing data analysis

BCL files produced by the Illumina sequencer were converted to FASTQ files using command mkfastq from the Cellranger analysis toolkit (v. 3.0.2 – Fig. 2, v 5.0.1 – Fig. 3, 10X Genomics). The FASTQ files were processed with Cellranger count to produce count matrices suitable for analysis in R using package Seurat^13^ (v. 4.0.3). Clustree^14^ was used to guide the selection of cluster resolution. The analysis script is available at <https://github.com/cemalley/Slamecka_methods>.

### Micro-RNA sequencing

The assay and a part of the data analysis was performed by Arraystar Inc.

RNA Quality Control

Agarose gel electrophoresis was used to check the integrality of total RNA samples. NanoDrop ND-1000 instrument was used for the measurement of concentration (abs 260) and protein contamination (ratio abs260/abs230) of total RNA samples.

Library Preparation

Reagents: NEBNext® Poly(A) mRNA Magnetic Isolation Module (New England Biolabs); RiboZero Magnetic Gold Kit (Human/Mouse/Rat) (Epicentre, an Illumina Company); NEBNext Small RNA Library Prep Set for Illumina (E7330L, New England Biolabs).

Total RNA of each sample was used to prepare the miRNA sequencing library, which included the following

steps:

1) 3’-adaptor ligation

2) 5’-adaptor ligation

3) cDNA synthesis

4) PCR amplification

5) size selection of 135 - 155 bp PCR amplified fragments (corresponding to 15 – 35 nt small RNAs). The libraries were denatured as single-stranded DNA molecules, captured on Illumina flow cells, amplified in situ as clusters, and finally sequenced for 51 cycles on Illumina NextSeq per the manufacturer’s instructions.

Sequencing

The DNA fragments in well mixed libraries were denatured to generate single-stranded DNA molecules, loaded onto channels of the flow cell at a concentration of 8 pM, and amplified in situ using TruSeq Rapid SR Cluster Kit (#GD-402-4001, Illumina). Sequencing was carried out using the Illumina NextSeq 500 according to the manufacturer’s instructions. Sequencing was carried out by running 51 cycles.

Sequencing Quality Control

Raw data files in FASTQ format were generated from the Illumina sequencer. To examine the sequencing quality, the quality score plot of each sample was plotted. Quality score Q is logarithmically related to the base calling error probability (P):

$$Q=-10\log_{10} P$$

miRNA-seq data analysis workflow

Raw sequencing data generated from Illumina NextSeq 500 that pass the Illumina chastity filter are used for following analysis. Trimmed reads (trimmed 3’-adaptor bases) were aligned to reference genome.

Quality Assessment of Sequencing Library

Agilent 2100 Bioanalyzer was used for assessment of the quality of sequencing library. Library concentration was determined by qPCR method.

Mapping Summary

After quality control, the reads were 3’-adaptor trimmed and filtered ≤ 15 bp reads with cutadapt software. The trimmed reads were aligned to reference genome with bowtie software. The reads statistical information is listed in the table below. In a typical experiment, it is possible to align 40-90% of the reads to the reference genome. However, this percentage depends on multiple factors, including sample quality, library quality, and sequencing quality.

miRNA Expression Results

The expression level (Reads count) of miRNA were calculated using miRDeep2^15^. The number of identified miRNA per group was calculated based on the mean of CPM in group ≥ 1. Counts per million reads (CPM) is calculated with the formula:

$$CPM=\frac{C \times{10}^{6}}{N}$$

C: The count of reads that map to a certain gene/transcript.

N: The total reads count that map to all genes/transcript.

Additional miRNA-seq analysis steps performed internally

The counts were processed in R (version 4.0.3) using package edgeR (version 3.34.0). Normalization, MDS plot construction, and differential expression analysis were performed using limma (version 3.48.1). Heatmaps were constructed using ComplexHeatmap (version 2.8.0).

### Methylated DNA immunoprecipitation sequencing

The assay and a part of the data analysis was performed by Active Motif.

Description of analysis steps performed by Active Motif

1. Sequence Analysis: The 75-nt single-end (SE75) sequence reads generated by Illumina sequencing (using NextSeq 500) were mapped to the genome using the BWA algorithm (“bwa aln/samse” with default settings). Alignment information for each read was stored in the BAM format. Only reads that passed Illumina’s purity filter, aligned with no more than 2 mismatches, and mapped uniquely to the genome were used in the subsequent analysis. In addition, duplicate reads were removed.

2. Determination of Fragment Density: Since the 5´-ends of the aligned reads (“tags”) represent the end of ChIP/IP-fragments, the tags were extended in silico (using Active Motif software) at their 3´-ends to a length of 200 bp, which corresponded to the average fragment length in the size-selected library. To identify the density of fragments (extended tags) along the genome, the genome was divided into 32-nt bins and the number of fragments in each bin was determined. This information (“signal map”; histogram of fragment densities) was stored in a bigWig file, which could be visualized in genome browsers. bigWig files also provided the peak metrics in the Active Motif analysis program described below.

3. Peak Finding: The generic term “Interval” is used to describe genomic regions with local enrichments in tag numbers. Intervals were defined by the chromosome number and a start and end coordinate. The two main peak callers used were MACS/MACS2^16^ and SICER^17^. MACS is suitable to identify the binding sites of transcription factors that bind to discrete sites (often containing a consensus DNA sequence) as well as many active histone marks and methyl-C enriched regions, while SICER is used to study proteins that bind to extended regions in the genome (such as repressive histone marks or RNA polymerase II). Both methods look for significant enrichments in the ChIP/IP data file when compared to the Input data file or relative to neighboring background regions.

4. Additional Analysis Steps - Normalization: The tag number of all samples (within a comparison group) was reduced by random sampling to the number of tags present in the smallest sample. This normalization method works well for most assays where the majority of tags map to background (i.e., non-peak) regions, and it can detect site-specific as well as global differences in target enrichments between samples.

5. Merged Region Analysis: To compare peak metrics between 2 or more samples, overlapping Intervals were grouped into “Merged Regions”, which are defined by the start coordinate of the most upstream Interval and the end coordinate of the most downstream Interval (union of overlapping Intervals; “merged peaks”). In locations where only one sample has an Interval, this Interval defines the Merged Region. The use of Merged Regions is necessary because the locations and lengths of Intervals are rarely the same when comparing different samples. Furthermore, with this approach fragment density values can be obtained even for samples for which no peak was called.

6. Annotations: After defining the Intervals and Merged Regions, their genomic locations along with their proximities to gene annotations and other genomic features were determined. In addition, average and peak (i.e., at “summit”) fragment densities within Intervals and Merged Regions were compiled.

MeDIP-seq analysis software versions used at Active Motif

bcl2fastq2 (v2.20): processing of Illumina base-call data and demultiplexing.

bwa^18^ (v0.7.12): alignment of reads to reference genome.

Samtools^19,20^ (v0.1.19): processing of BAM files.

BEDtools^21^ (v2.25.0): processing of BED files.

MACS2 (v2.1.0): peak calling; narrow peaks.

SICER (v1.1): peak calling: broad peaks.

wigToBigWig^22^ (v4): generation of bigWIG files.

Additional MeDIP-seq analysis steps performed internally

Modified differential peak analysis

To conservatively assess differential methylation, only those peaks supported by at least two biological replicates were retained for downstream analysis using basic filtering and chaining with dplyr (tidyverse^23^ v1.3.1) in R. DESeq-2^24^ (v1.42.0) with default parameters was then used to perform differential methylation analysis to which the filtered MeDIP-Seq normalized peak counts were supplied.

Manhattan plots were generated in ggplot2^8^ (v3.3.5). Top 20 significantly differentially methylated peaks were highlighted.

Heatmaps were created using pheatmap^25^ (v1.0.12) by plotting the unscaled log_2_(count + 1) for the top 50 peaks sorted by the absolute value of the log_2_(fold-change). We annotated the sample type using the options for annotation_col and annotation_colors in pheatmap.

Locus zoom plots were built using Gviz (v 1.36.2) by creating and combing ideogram tracks, biomaRt (v2.48.2) generated annotation tracks, custom annotation tracks for identified MeDIP-Seq peaks, gene axis tracks and count data tracks highlighting approximate promoter regions (1000 bp upstream of the transcriptional start site), and significant differentially methylated peaks within the specified locus.

Count data for long arm of chromosome 21 was extracted from BAM files for each sample and replicate. Then the data was split into 40 numbered bins and a histogram was generated in ggplot2 (v3.3.5).

25.	Kolde, R. pheatmap: Pretty Heatmaps. https://CRAN.R-project.org/package=pheatmap.
